# INCREASED CHLOROPLAST OCCUPANCY IN BUNDLE SHEATH CELLS OF RICE *hap3H* MUTANTS REVEALED BY CHLORO-COUNT, A NEW DEEP LEARNING-BASED TOOL

**DOI:** 10.1101/2024.06.23.600271

**Authors:** Julia Lambret-Frotte, Pedro P. Buarque de Gusmão, Georgia Smith, Shuen-Fang Lo, Su-May Yu, Ross W. Hendron, Steven Kelly, Jane A. Langdale

## Abstract

There is an increasing demand to boost photosynthesis in rice to increase yield potential. Chloroplasts are the site of photosynthesis, and increasing the number and size of these organelles in the in leaf is a potential route to elevate leaf-level photosynthetic activity. Notably, bundle sheath cells do not make a significant contribution to overall carbon fixation in rice and thus various attempts are being made to increase chloroplast content in this cell type. In this study we developed and applied a deep learning tool named Chloro-Count to demonstrate that loss of *OsHAP3H* function in rice increases chloroplast occupancy in bundle sheath cells by 50%. Although limited to a single season, when grown in the field *Oshap3H* mutants exhibited increased numbers of tillers and panicles as compared to controls or gain of function mutants. The implementation of Chloro-Count enabled precise quantification of chloroplasts in loss- and gain-of-function *OsHAP3H* mutants and facilitated a comparison between 2D and 3D quantification methods. In wild-type rice, as the dimensions of bundle sheath cells increase, the volume of individual chloroplasts also increases. However, the larger the chloroplasts the fewer there are per bundle sheath cell. This observation revealed that a mechanism operates in bundle sheath cells to restrict chloroplast occupancy as cell dimensions increase. That mechanism is unperturbed in *Oshap3H* mutants. The use of Chloro-Count also revealed that 2D quantification, upon which most previous studies have relied, is compromised by the positioning of chloroplasts within the cell. Chloro-Count is therefore a valuable tool for accurate and high-throughput quantification of chloroplasts that has enabled the robust characterization of *OsHAP3H* effects on chloroplast biogenesis in rice. Whereas previous studies have increased chloroplast occupancy in bundle sheath cells by increasing the size of individual chloroplasts, loss of *OsHAP3H* function leads to an increase in chloroplast numbers.

## INTRODUCTION

There is an urgent need to boost global crop production and photosynthesis is an important target for manipulation (Ort *et al*., 2015). In leaves of the majority of plant species, photosynthesis primarily occurs in mesophyll cells, with barely detectable contributions from chloroplasts in the bundle sheath cells that surround leaf veins. Increasing chloroplast function in bundle sheath cells of these ‘C_3_’ photosynthesizing species is thus one potential strategy to improve overall photosynthetic capacity in the leaf. To this end, a number of genetic targets for manipulation have been identified. However, the limited capacity to accurately quantify chloroplast occupancy in the leaf has thus far hampered evaluation of the phenotypic consequences of any such manipulations. Most quantifications of chloroplast occupancy to date have been carried out using two-dimensional (2D) images obtained via light or electron microscopy (Pyke *et al*., 1994; Kubínová *et al*., 2014; Khoshravesh *et al*., 2016; Lee *et al*., 2021; Plackett & Hibberd, 2024) but where comparisons have been made, estimations of total chloroplast volume from 2D sections were significantly lower than those calculated from 3D reconstructions (Harwood *et al*., 2020). Furthermore, in the specific case of bundle sheath cells, 2D assessments were misleading because chloroplast distribution within the cell is very heterogeneous (Williams *et al*., 1989; Harwood *et al*., 2020). Development of a high throughput method for quantifying chloroplast number and volume in 3D would thus increase the potential to identify plant lines with improved photosynthetic capacity.

3D reconstruction of cells can be achieved by a myriad of techniques, including serial block face scanning electron microscopy (SBF-SEM) and confocal laser scanning microscopy (CLSM). A comparative analysis that used both techniques to build 3D structures of bundle sheath cells in C_4_ plants showed that SBF-SEM was more reliable than CLSM (Lee *et al*., 2023) but the costly machinery and time-consuming sample preparation makes high-throughput analysis via SBF-SEM impractical in most cases. As such, an alternative quantitative approach that uses CLSM is needed. Artificial intelligence has increasingly found application in biological imaging for precise reconstruction of plant cells and organs (Fernandez *et al*., 2010; Barbier de Reuille *et al*., 2015; Wolny *et al*., 2020; Gómez-de-Mariscal *et al*., 2021; Vijayan *et al*., 2021) but there are limited tools available for chloroplast quantification in plant cells (Li *et al*., 2021; Feng *et al*., 2023; Su *et al*., 2023). A suitable method to quantify chloroplast number and volume in 3D would take advantage of chlorophyll autofluorescence so that minimal sample preparation was required (Billakurthi and Hibberd 2023) and would rapidly process thousands of serial images along the z-axis of the specimen. To the best of our knowledge, none of the existing neural networks trained for chloroplast quantification integrate instance segmentation and volume estimation with CLSM imaging (Li *et al*., 2021; Feng *et al*., 2023; Su *et al*., 2023).

Here we have manipulated the function of a gene that is predicted to regulate chloroplast development in rice and have characterized the phenotype of transgenic lines using a deep learning-based tool that we developed to quantify chloroplast number and volume. Specifically, we generated loss- and gain-of-function mutants in a rice *HEME-ASSOCIATED PROTEIN* (*OsHAP*) gene. The *HAP* genes, also known as NUCLEAR FACTOR Y (NF-Y) or CCAAT-binding factor (CBF) genes, comprise three subgroups (*HAP2/NF-YA/CBF-B*, *HAP3/NF-YB/CBF-A* and *HAP5/NF-YC/CBF-C*) which form heterotrimeric complexes that regulate gene expression across diverse biological processes (Petroni *et al*., 2012). Notably, silencing of *OsHAP3A* reduces chlorophyll accumulation and impairs chloroplast biogenesis in rice mesophyll cells (Miyoshi *et al*., 2003) and another of the *HAP3* genes (*OsHAP3H*) maps to the same locus as *OsCAR8* and *Ghd8* which were shown by QTL analysis to influence carbon assimilation rates via photosynthesis and grain productivity, respectively (Miyoshi et al. 2003, Thirumurugan et al. 2008, Yan et al. 2011, Adachi et al. 2017). We thus reasoned that targeted manipulation of *OsHAP3* could activate chloroplast development in bundle sheath cells. Using our newly developed tool, henceforth named Chloro-Count, we quantified chloroplast number and volume to reveal a significant increase in total chloroplast occupancy in bundle sheath cells of loss-of-function *Oshap3H* mutant lines.

## RESULTS

### *OsHAP3H* as a candidate for manipulation in rice

Members of the *HAP3* gene family have a histone-like transcription factor domain that mediates protein-protein interactions to form a heterotrimer that binds CCAACT DNA sequences (Figure 1A; Petroni et al. 2012). With a view to enhancing chloroplast development and photosynthetic capacity in rice, we first examined phylogenetic relationships within the gene family (Figure 1B). The phylogeny reveals that two distinct clades were present in the last common ancestor of rice, maize, wheat and Arabidopsis (HAP3A-C and HAP3D-K) and the topology of sub-clades indicates that all 11 rice genes were present prior to speciation. Through the analysis of antisense mutants, *OsHAP3A* has been shown to control chloroplast development, with *OsHAP3B* and *OsHAP3C* also possibly implicated (Miyoshi *et al*., 2003). By contrast, a role for *OsHAP3H* in photosynthesis has been inferred by QTL analysis. *OsHAP3H* maps to the same locus as *OsCAR8* and *Ghd8*, which were independently identified as QTLs that led to higher carbon assimilation and increased grain yield respectively (Thirumurugan *et al*., 2008; Yan *et al*., 2011; Adachi *et al*., 2017). Specifically, a nonsense mutation in *OsCAR8* (Habataki) resulted in a truncated protein and led to higher carbon assimilation as compared to *OsCAR8* (Hoshihikari) (Figure 1A; Adachi et al. 2017). Similarly, a SNP in the start codon of *Ghd8* (Zhenshan) caused a frameshift that resulted in a premature stop codon and when two functional alleles [*Ghd8* (93-11) and *Ghd8* (Nipponbare)] were independently introduced into the Zhenshan (ZS) background, plants showed 60% more grain yield than their ZS counterparts (Figure 1A; Yan et al. 2011). Given the inter-relatedness between chloroplast biogenesis, photosynthetic carbon assimilation and grain yield, we decided to directly test the role of *OsHAP3H* in chloroplast development through gain and loss of function analyses in a single rice variety.

**Figure 1.**
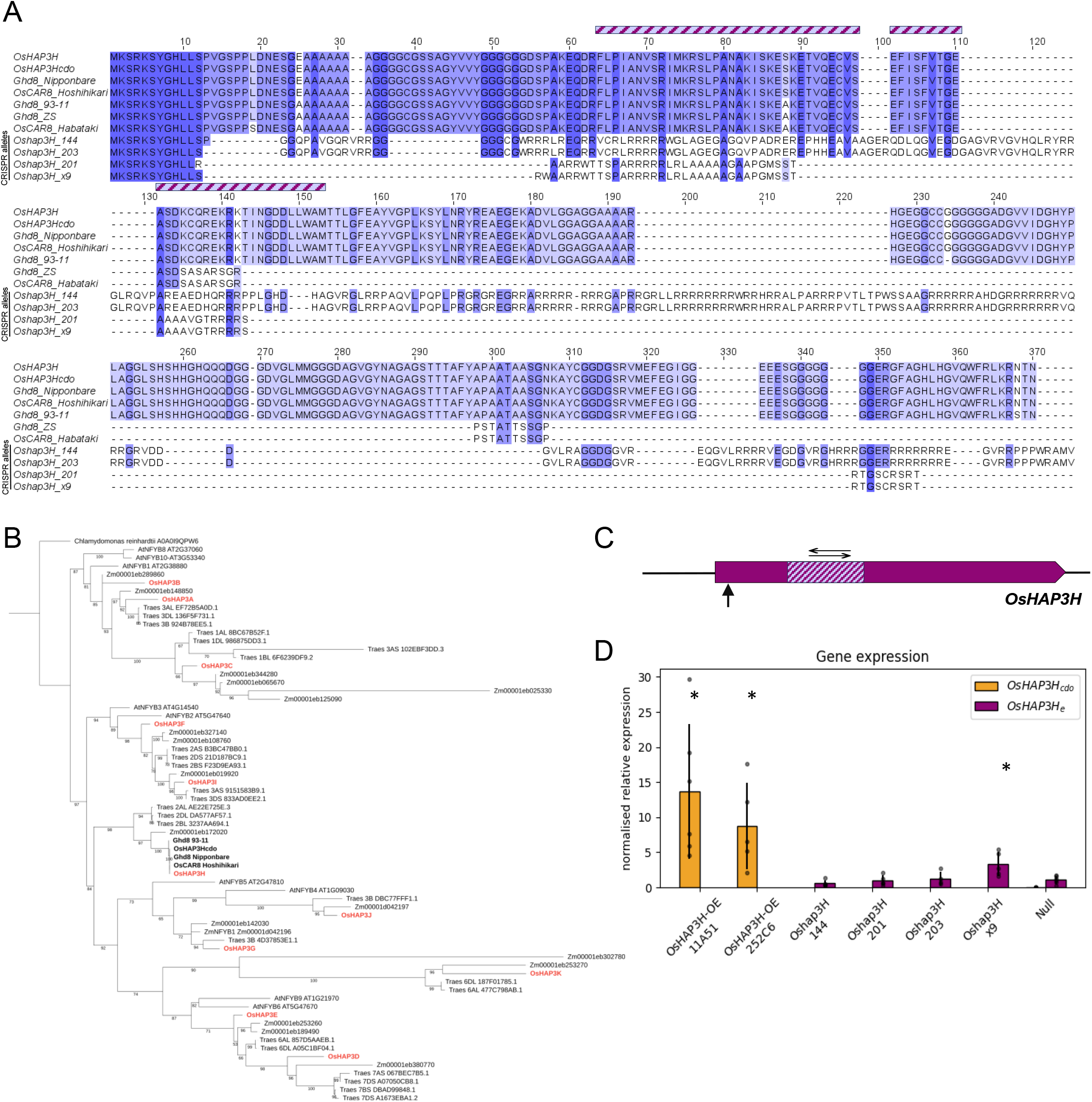
Mutant alleles of *OsHAP3H*. **A)** Protein sequence alignment showing conserved regions between *OsHAP3H*, previously reported *OsCAR8* and *Ghd8* alleles and mutant *Oshap3H* alleles generated in this study. Textured bar shows the histone-like transcription factor domain present in HAP3/NF-YB/CBF-A proteins. **B)** Phylogenetic analysis of the HAP3 family was inferred by Maximum Likelihood and evolutionary distances were computed using the JTT+F+I+G4 method. Clade support was calculated by 1000 bootstrap replicates. **C)** Gene model of *OsHAP3H* where purple bar indicates coding sequence flanked by untranslated regions depicted as black lines. The hatched region depicts the histone-like transcription factor domain characteristic of HAP3 proteins, the vertical arrow shows the position of the gRNA used to create *Oshap3H* mutants and the horizontal arrows indicate the amplicon assessed for gene expression analysis. **D)** Quantitative analysis of gene expression via qPCR in *OsHAP3H-OE*, *Oshap3h* and null segregating lines. Relative normalized expression of *OsHAP3H*_cdo_ and *OsHAP3H*_e_ was calculated based on the expression of two reference genes for each sample. Each dot represents a biological replicate. Statistical significance was calculated using a Student’s t-test and pairwise comparison with the null segregant sample. Asterisks show statistical different groups compared to the null segregant.

To test the hypothesis that manipulation of *OsHAP3H* could enhance chloroplast development in rice, gain- and loss-of-function mutants were generated in *Oryza sativa spp japonica* cv Kitaake. For gain of function, the coding region of *OsHAP3H* was expressed under the control of the constitutive maize ubiquitin promoter (*ZmUBI*_pro_; Figure S1A). To prevent potential gene silencing of the native coding sequence via miRNA activity and to facilitate the differentiation of transcripts originating from the overexpression construct from those originating from the native gene, multiple silent mutations were introduced along the coding sequence (*OsHAP3H_cdo_*; Figure 1A, Figure S1B). Two *OsHAP3H_cdo_*-OE lines were generated, each containing two independent T-DNA insertions, along with a null segregant line (Figure S1C). For loss of function, CRISPR/Cas9-directed mutagenesis was carried out using a single guide RNA targeting the only exon, and four independent *Oshap3H* lines harbouring different nonsense mutations were generated (Figure 1A, Figure S1D). An assessment of predicted protein sequences revealed that in all four mutant alleles the histone-like transcription factor domain is disrupted, impairing the ability to form the protein-protein interactions that are necessary for regulating the transcription of downstream targets (Figure 1A, C). Gene expression analysis revealed that, although variable, *OsHAP3H_cdo_* transcript levels were, on average, 10 times higher in *OsHAP3H_cdo_*-OE lines than in the null segregant (Figure 1D). As expected, in most of the *Oshap3H* lines, levels of endogenous (but nonsense mutated) *OsHAP3H* transcripts (*OsHAP3H_e_*) were the same as in the null line (Figure 1D). In one loss of function line, however, *OsHAP3H_e_* levels were five times higher than in the null, a level comparable to that seen in some of the *OsHAP3H_cdo_*-OE samples (Figure 1D). The reason for this anomaly is not known, but given it was only seen in one out of four lines it cannot be attributed to loss of *OsHAP3H* function. Together therefore, two gain- and four loss-of-function lines had been generated for phenotypic analysis of chloroplast and photosynthetic parameters.

### Loss of function *Oshap3H* mutants accumulate higher chlorophyll levels than wild-type and exhibit enhanced photosynthetic activity in the field

To determine whether manipulation of *OsHAP3* activity had any impact on photosynthesis, we first measured chlorophyll levels in gain- and loss- of function lines grown in the growth chamber. Notably the accumulation of both chlorophyll a and b were significantly higher in *Oshap3H* lines than in *OsHAP3H_cdo_-*OE or null groups (Figure 2A, B). To test whether the elevated chlorophyll levels in loss-of-function *Oshap3H* mutants provided any physiological benefit in a field context, lines were grown in randomised plots in the GM Experiment Station in NCHU, Wufeng District, Taichung City, Taiwan from August to November 2022. Because null segregants could not be obtained from the CRISPR/Cas9 experiment (all regenerated lines were mutant and the Cas9 could not be segregated away by crossing), a null segregant from the overexpression experiment was selected randomly to be used as a tissue-culture derived control and non-transformed Kitaake lines were used as a second control. Several growth, environmental and photosynthetic parameters were measured (Figure 2C-O). Plant height did not differ between any of the analysed lines (Figure 2C). However, *Oshap3H* lines developed significantly more tillers as compared to null and Kitaake lines (Figure 2D). Despite this increase, mutants did not accumulate significantly more total biomass, above ground biomass or show any increase in yield (Figure 2E-G). Surprisingly, unlike seen with growth chamber grown plants, *Oshap3H* plants did not accumulate more chlorophyll than null and Kitaake lines when grown in the field (Figure 2H). Similarly, although panicle number was significantly increased (Figure 2I), neither weight or length differed from controls (Figure 2J, K). Despite the increase in both panicle number and tiller number, *Oshap3H* lines did not exhibit higher yield possibly due to slight reduction in seed weight compared to null and Kitaake (Figure 2L). However, under similar photosynthetically active radiation (PAR), relative electron transport rate (ETR) and the quantum efficiency of photosystem II (LJPSII) were significantly higher in *Oshap3H* lines than in null or Kitaake lines (Figure 2M-O). With the caveat that these data were collected from just a single field season, the observations reported suggest that loss of *OsHAP3H* function can enhance photosynthetic activity in rice.

**Figure 2.**
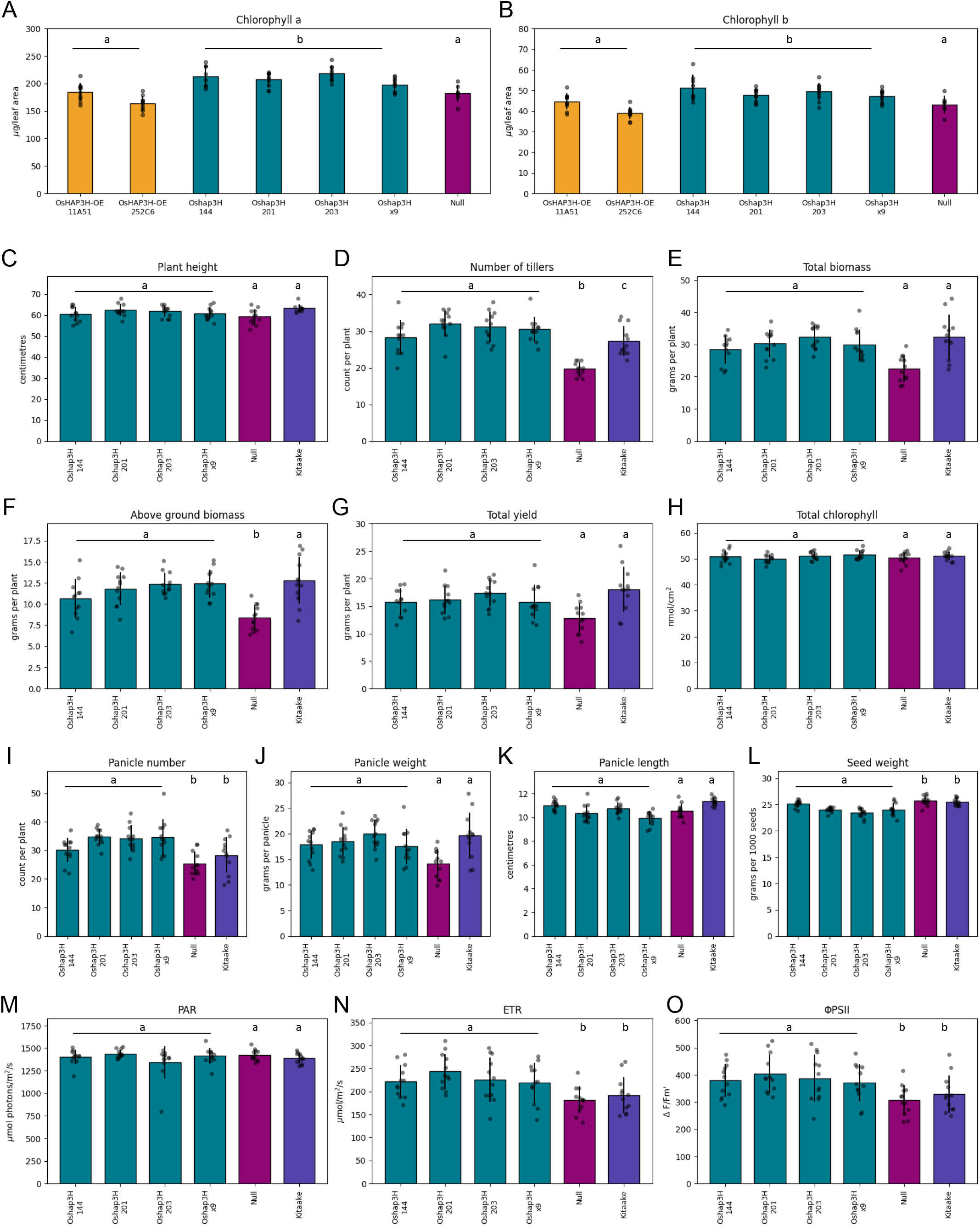
Loss of function *Oshap3H* lines exhibit enhanced photosynthetic capacity. **A-B)** Quantification of chlorophyll a (A) and chlorophyll b (B) from *OsHAP3H*-OE, *Oshap3H* and null segregant plants grown in a growth chamber. **C-O)** *Oshap3H* lines grown in the field were assessed for plant height (C); number of tillers (D); total biomass (E); above ground biomass without panicles (F); total yield (G); total chlorophyll (H); panicle number (I), weight (J) and length (K); seed weight (L); photosynthetically active radiation (PAR; M); relative electron transport rate (ETR; N); and quantum efficiency of photosynthesis (LJPSII, O). Each dot represents one plant, the coloured bars are the mean value and the black lines are the associated standard deviations. Statistical significance was calculated using ANOVA and a Tukey HSD test based on untransformed data. Lines from the same genetic background were grouped together and treated as one statistical unit. Different letters indicate statistically different groups (p-value ≤0.05). See Table S1 for raw data.

### Development of a deep learning-based tool to quantify chloroplast volume in rice leaves

Having established that loss of *OsHAP3H* function can result in perturbed chlorophyll and/or photosynthetic parameters in rice, we next sought to determine the impact on chloroplast development. In particular, we wanted to know the extent to which the mutant phenotype was associated with enhanced chloroplast development in bundle sheath cells because chloroplast development is normally limited in this cell-type. The quantification of chloroplast volume in bundle sheath cells involves the analysis of sequential images that span all cell dimensions. To increase the scale and accuracy of chloroplast measurements we developed Chloro-Count, a tool based on deep learning, to semi-automate the analysis of a large number of images. To prepare cells for imaging, segments around intermediate veins of rice leaves were first isolated and digested to release mesophyll cells from the vascular strand, and then physically agitated to release bundle sheath cells (see Methods). Sequential z-stack images of single bundle sheath cells were captured using CLSM (Figure 3A) and then chloroplasts and bundle sheath cells were manually segmented and labelled, with each chloroplast receiving a unique label within a bundle sheath cell. The annotated images were partitioned into train, validation, and test sets that were used to ground-truth two distinct Chloro- Count models that separately identified individual chloroplast and bundle sheath cell areas across the z-stacks. For chloroplast area, the model test showed that Chloro-Count produced an Average Recall (AR) of 0.62 and an Average Precision (AP) of 0.89, i.e. Chloro-Count was able to retrieve over 62% of all manually annotated chloroplast regions and 89% of the retrieved segments mapped to true chloroplasts, with an Intersection-over-Union (IoU) value of at least 0.5 (Figure 3B). For bundle sheath cell area, the model performed even better, with an AR of 0.768 and an AP of 0.927. To increase accuracy, a manual verification step of segmentation was performed for every cell to eliminate any false negatives, prior to cell and chloroplast volume calculations. Volumes were calculated by multiplying the area of the object in each z-stack by the height of the stack, and then summing the values within and between all of the z-stacks that the object occupied (Figure 3A). Crucially, output from this deep learning-based tool facilitated high-throughput quantification of chloroplast and bundle sheath cell volume in rice leaves, providing the opportunity for robust comparisons between 2D and 3D quantifications, and between wild-type and mutant leaves.

**Figure 3.**
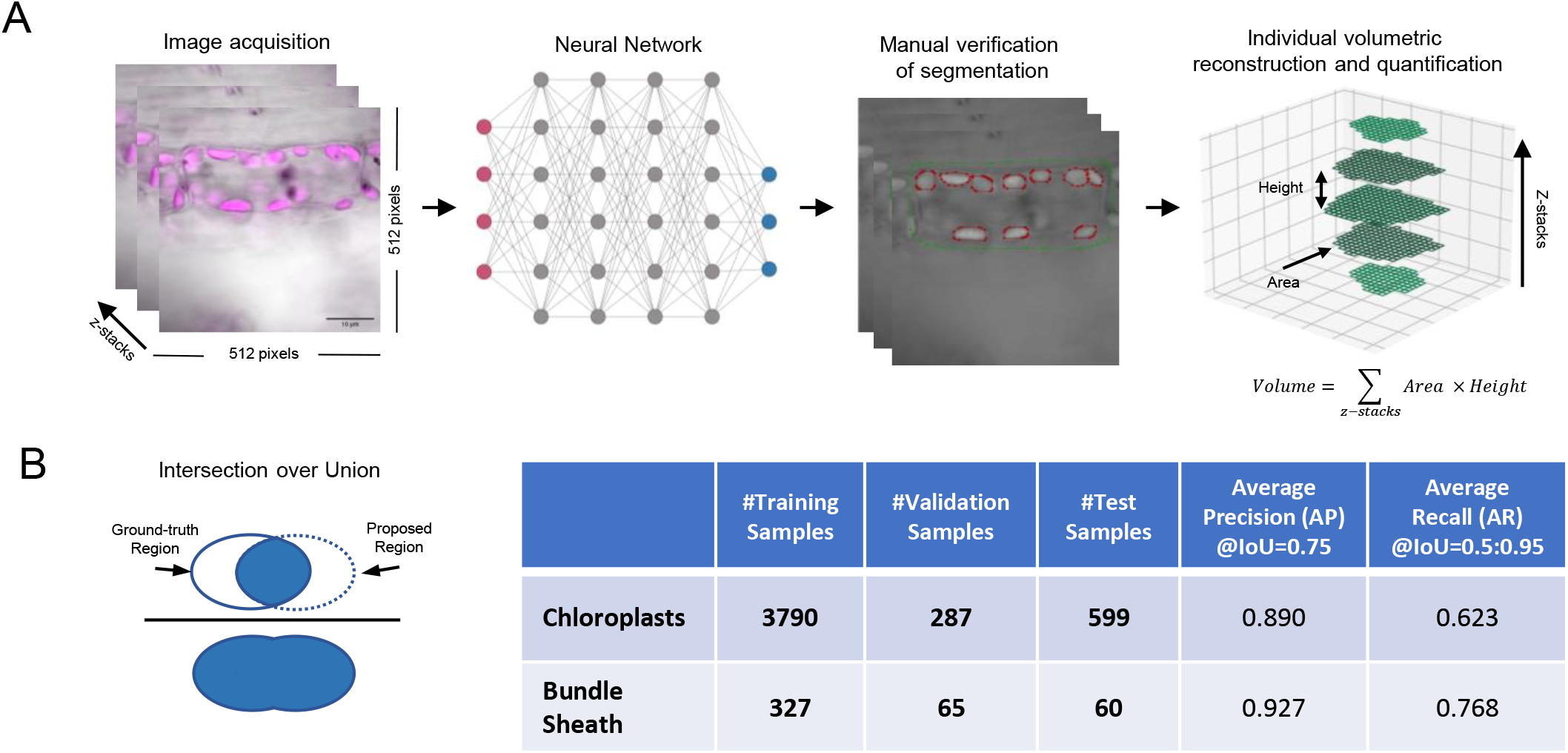
Overview of Chloro-Count. **A)** Flowchart of image processing and data analysis for measuring chloroplast and bundle sheath cell volumes. Bundle sheath cell images were taken with CLSM to span all cell dimensions. Images were used to train a neural network model and, after manual verification of segmentation by the model, individual volume reconstruction of chloroplasts and bundle sheath cells was performed by summing the product of the segmented area times the distance between z-stacks. **B)** The diagram shows the Intersection of Union (IoU) that compares the overlap between the ground truth region of the chloroplast and its proposed region identified by the Chloro-Count model. The table shows the number of chloroplast and bundle sheath samples used to train, validate and test the Chloro-Count model. It also shows model performance measured by Average Recall (AR) and Average Precision (AP) which indicate, respectively, the ability of the model to correctly detect an object (AR) and how many of the objects were correctly identified (AP), with an IoU of at least 0.5.

### Cell and chloroplast dimensions are poorly estimated on the basis of area

Comparisons between 2D and 3D estimates of bundle sheath and chloroplast dimensions were performed on leaves of wild-type Kitaake rice plants, using Chloro-Count to obtain 3D estimates (bundle sheath volume, average chloroplast volume, chloroplast count, and bounding box dimensions). Harwood et al. (2020) determined that the volume of C_3_ grass chloroplasts is, on average, 26.8±7.3 µm^3^. To eliminate potential image artifacts mistakenly annotated as chloroplasts, objects with a volume estimate of less than half this average (13.4 µm³) were excluded (Harwood *et al*., 2020). 2D estimates (bundle sheath area, average chloroplast area, and chloroplast count) were retrieved for each sample from the z-stack with the highest chloroplast count and were compared to 3D measurements by linear regression (Figure 4, Table S1). Notably, because the area of a longitudinal section (z-stack) does not account for variations in either the length or diameter of a cell (both of which are important given bundle sheath cells are cylindrical in shape), correlations between 2D and 3D estimates of cell dimensions are poor (R^2^ = 0.0019, p-value = 0.785; Figure 4A). However, the ratio of x/y axes from the bounding boxes of a cell are a reasonable proxy of cell volume, given the statistically significant correlation observed (R^2^ = 0.2387, p-value = 0.0010; Figure S2). Average chloroplast area also shows weak correlation with corresponding average volumes (R^2^ = 0.0435, p-value = 0.1849) (Figure 4B), but chloroplast numbers obtained from 2D and 3D estimates show a statistically significant correlation (R^2^ = 0.6214, p-value = 5.7 x 10^-10^) (Figure 4C). Because chloroplasts are ovoid or flat concave disc structures (Harwood *et al*., 2020), a single longitudinal section possibly captures part of most chloroplasts present, but it will not accurately represent actual sizes due to diverse orientations within the cell. Moreover, chloroplast distribution around the cell is not uniform; distribution across the proximal-distal axis of the cell is relatively consistent when compared to the medio-lateral axis, where chloroplasts are preferentially localized towards the edges of the cell (Figure S3). These observations are consistent with the idea that the size and position of the vacuole has a greater impact on chloroplast density measurements when viewed in the medio- lateral and adaxial-abaxial leaf axes than in the proximo-distal leaf axes. Nevertheless, the relative content of chloroplasts estimated in 2D and 3D are significantly correlated (R^2^ = 0.5154, p-value = 8.7 x 10^-8^, Figure 4D). In summary, whereas chloroplast numbers and relative occupancy in a cell can be quantified equally well via 2D (particularly in paradermal/longitudinal sections) or 3D approaches, estimating the real dimensions of bundle sheath cells and the chloroplasts within them needs 3D analysis.

**Figure 4.**
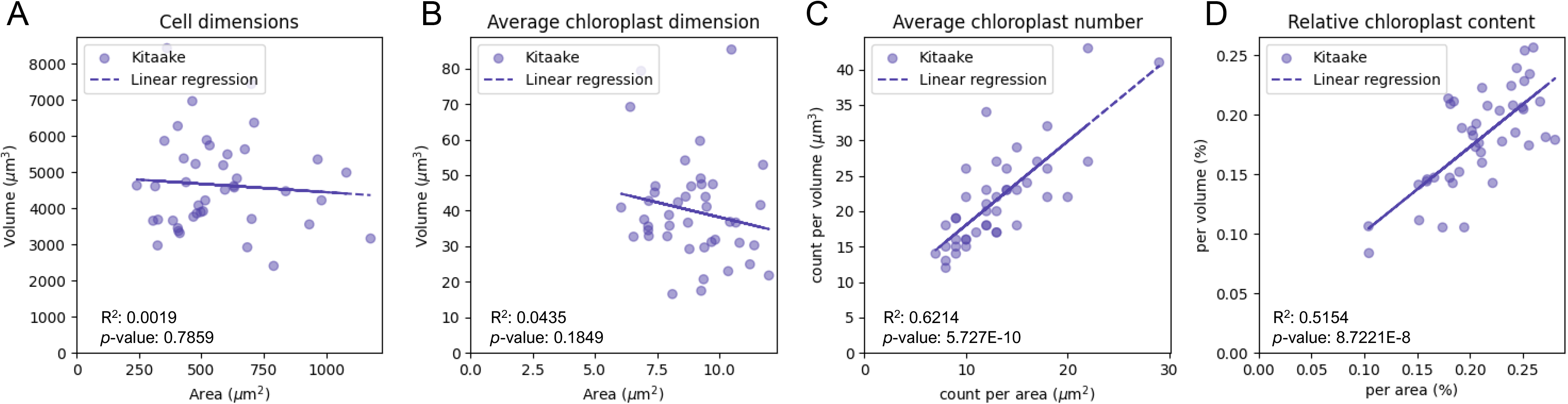
Comparison between 2D and 3D measurements of chloroplasts and bundle sheath cells in rice. **A)** Scatterplot of bundle sheath cell area measured on a single z-stack with the highest number of chloroplasts compared to the volume of the same cells measured via Chloro-Count. **B)** Average chloroplast area per cell measured on a single z-stack compared to the average volume of chloroplasts in a bundle sheath cell. **C)** Average chloroplast number measured on a single z-stack compared to the total chloroplast number measured in the total volume of a bundle sheath cell. **D)** Relative chloroplast content calculated by dividing the sum of all chloroplasts in a single z-stack to the area of the bundle sheath cell in the same stack; plotted against the summed volume of all chloroplasts in a cell divided by the volume of the cell. The measurements are shown in percentages. Solid lines show linear regression of data obeying a normal distribution; R^2^ and p-values show variation and significance of the model for each analysis, respectively. See Table S1 for raw data.

### Chloroplast occupancy in bundle sheath cells of *Oshap3H* mutants is increased by ∼50%

Having established that accurate measurements of chloroplast content in bundle sheath cells can only be achieved via 3D approaches, we used Chloro-Count to analyse *OsHAP3H* mutants. Analysis of variance revealed that both chloroplast and bundle sheath measurements varied significantly between samples in the same line (Table S1). For statistical purposes, to minimise variation effects, lines of either gain- or loss-of-function mutants were therefore combined in classes and treated as one statistical entity (Table S1). In this context, no statistical difference was detected in bundle sheath cell volume between any of the analysed groups (i.e. gain of function vs loss of function vs null) (Figure 5A). Similarly, there was no significant difference in average chloroplast volume between *OsHAP3H_cdo_*-OE, *Oshap3H* and null lines (Figure 5B). As such, neither gain or loss of *OsHAP3H* function affects bundle sheath cell or chloroplast volume. By contrast, *Oshap3H* lines showed ∼30% more chloroplasts per bundle sheath cell than either *OsHAP3H_cdo_*-OE or null lines, which were not statistically different from each other (Figure 5C). Importantly, the increase in chloroplast number in *Oshap3H* lines led to a 50% increase in relative chloroplast occupancy in bundle sheath cells (Figure 5D). In *Oshap3H* lines 144 and 203, chloroplasts occupied 14% (on average) of the bundle sheath cell volume, whereas this number was 11% for lines 201 and x9. Both of these numbers are significantly higher than the 8% seen in the *OsHAP3H_cdo_*-OE and null lines (Figure 5D). Collectively these data reveal that loss of *OsHAP3H* function increases the number of chloroplasts in bundle sheath cells of rice such that chloroplast occupancy in that cell type is enhanced by fifty per cent.

**Figure 5.**
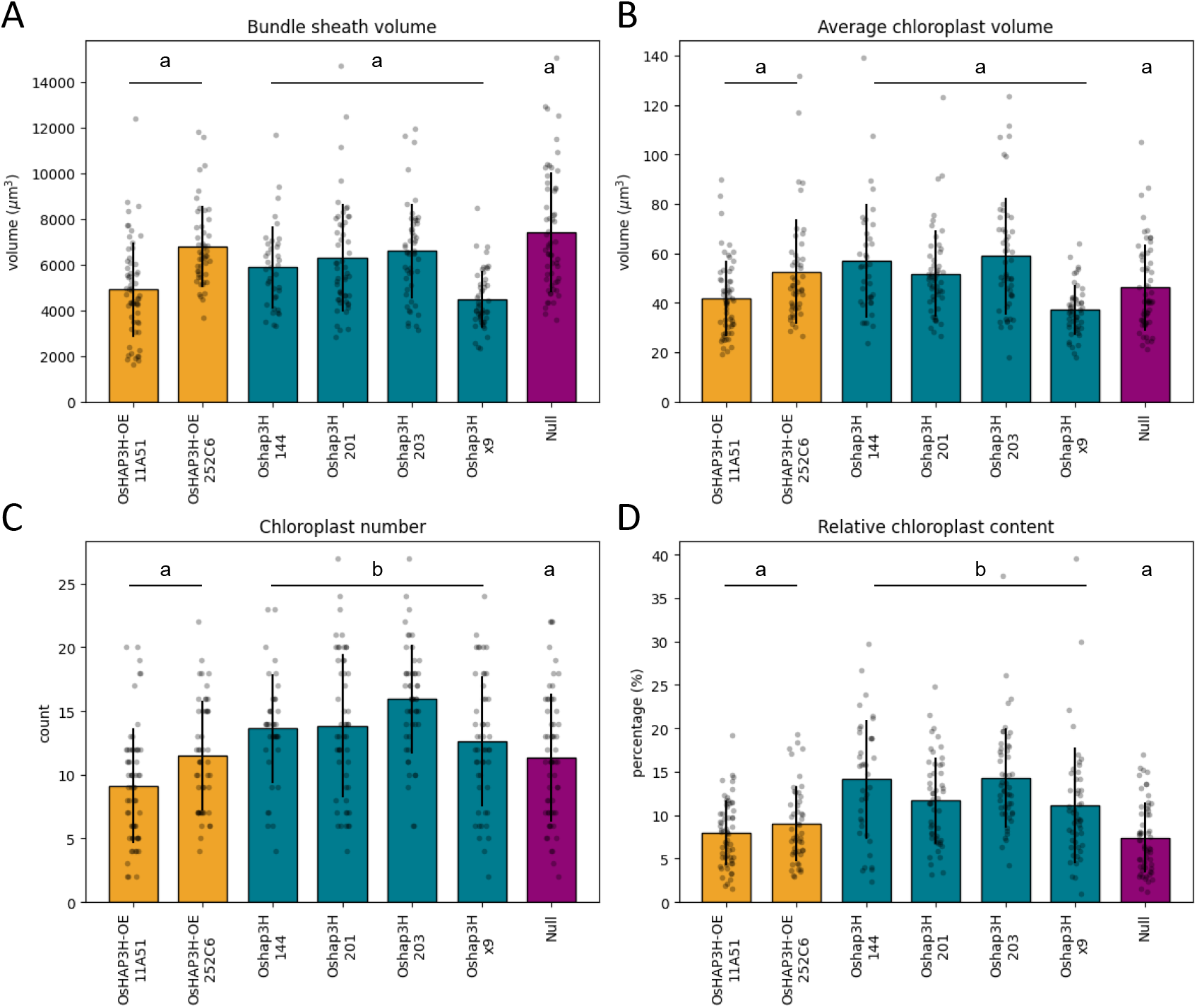
Chloroplast volume in bundle sheath cells is increased in *Oshap3H* mutants. **A-D)** Quantification of bundle sheath cell volume (A), average chloroplast volume (B), chloroplast number (C) and relative chloroplast content (D). Chloroplast quantifications were calculated per bundle sheath cell and each dot represents the data from a single cell. An average of 10 cells from a minimum of four plants was assessed for each line. Statistical significance was calculated using ANOVA and a Tukey HSD test based on data that was transformed to a normal distribution. Different letters indicate statistically different groups (p-value ≤0.05). Lines from the same genetic background were grouped together and treated as one statistical unit. See Table S1 for raw data.

### Constraints on increased chloroplast occupancy in rice bundle sheath cells revealed through 3D analysis

Given that the 50% increased chloroplast occupancy in *Oshap3H* mutant lines led to modest increases in chlorophyll accumulation (in growth chamber condition), electron transport rate and operating efficiency of PSII (under field condition), we next sought to determine what parameters might be constraining further increases in chloroplast occupancy in bundle sheath cells. Linear regression analysis of WT Kitaake rice plants revealed significant correlations both between average chloroplast area and cell area (R^2^ = 0.2452, p-value = 0.0008) and between average chloroplast volume and cell volume (R^2^ = 0.4847, p = 3.0 x 10^-7^), but average chloroplast volume was better explained by cell volume (R^2^ = 0.48), than average chloroplast area was by cell area (R^2^ = 0.24; Figure 6A, B). By contrast, the correlation between chloroplast number and cell area was statistically significant (R^2^ = 0.5651, p-value = 9.5 x 10^-9^), whereas there was no correlation between chloroplast number and cell volume (R^2^ = 0.0043, p-value = 0.6775; Figure 6C, D). As such, our data suggest that chloroplast number does not increase as cell volume increases, but chloroplast volume does. Further examination of the relationship between chloroplast number and average chloroplast dimensions revealed that in 2D measurements, there is no statistically significant correlation between average chloroplast area and chloroplast number (R^2^ = 0.0453, p-value = 0.1759; Figure 6E), but in 3D estimates, a statistically significant negative correlation exists between average chloroplast volume and chloroplast number (R^2^ = 0.1033, p-value = 0.0378; Figure 6F). These observations suggest that bundle sheath cell volume in rice constrains the total chloroplast occupancy by balancing both the size and number of individual chloroplasts.

**Figure 6.**
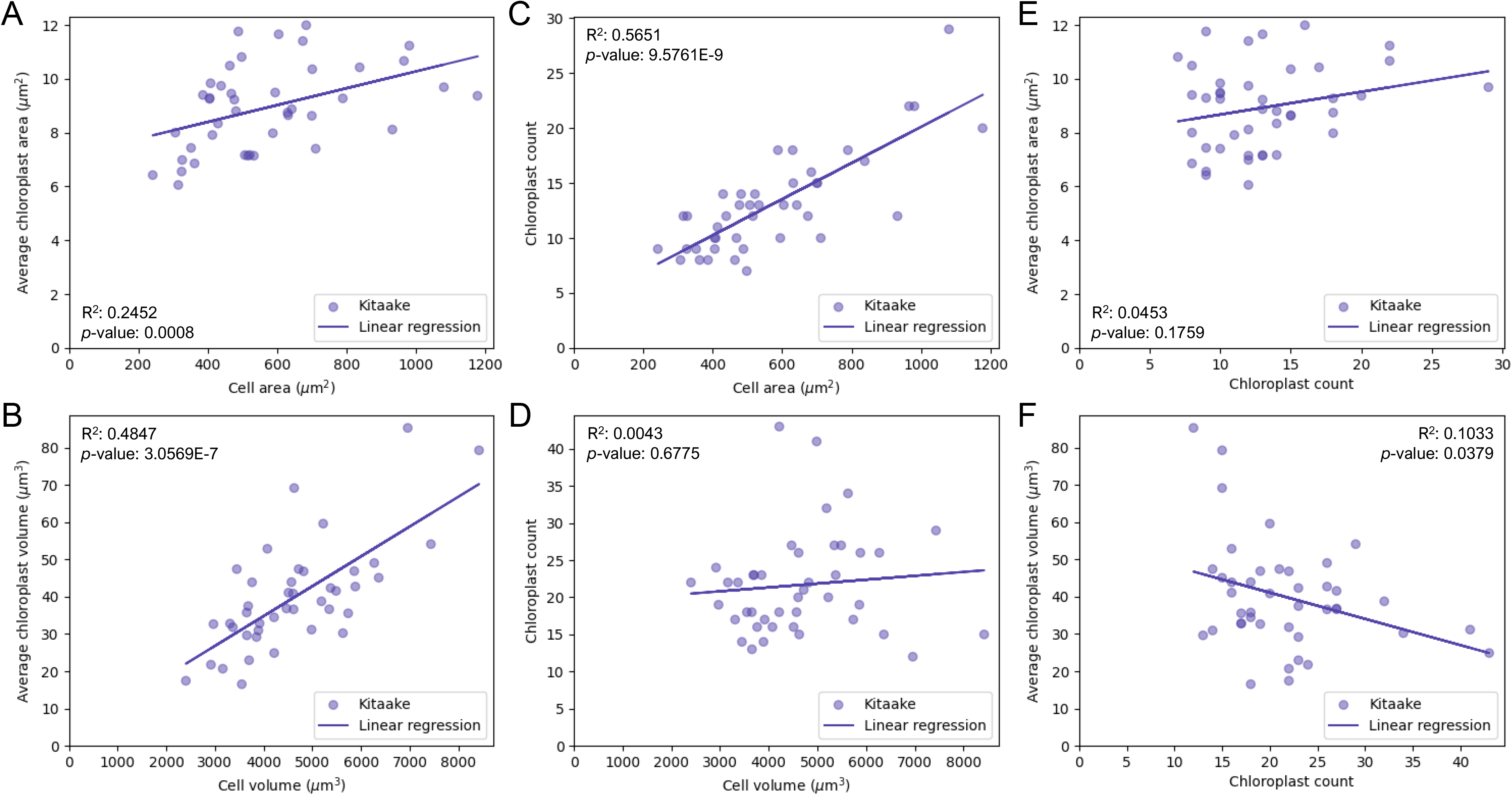
Bundle sheath cell volume constrains average chloroplast number. **A, B)** Effect of bundle sheath cell area on average chloroplast area (A) and bundle sheath cell volume on average chloroplast volume (B). **C, D)** Effect of bundle sheath cell area on average chloroplast number (C) and bundle sheath cell volume on average chloroplast number (D). **E)** Correlation between chloroplast number in one z-stack and average chloroplast area. **F)** Correlation between total chloroplast number and average chloroplast volume. Solid lines show linear regression of data following a normal distribution; R^2^ and p-values show variation and significance of the model for each analysis, respectively. See Table S1 for raw data.

To evaluate whether bundle sheath cell volume could be restricting any further increase in chloroplast occupancy above 50% in *Oshap3H* mutant lines, we measured the volume and number of chloroplasts in individual bundle sheath cells of *OsHAP3H-OE,* null and *Oshap3H* lines. *Oshap3H* lines show a positive correlation between chloroplast number and average chloroplast volume, with borderline statistical support (R^2^ = 0.1508, p-value = 0.037; Figure 7). However, analysis of covariance did not show any statistical significance between *Oshap3H*, null or *OsHAP3H-OE* lines (p-value = 0.7178). Therefore, although loss of *OsHAP3H* function led to an increase in chloroplast number in bundle sheath cells, the level of that increase was likely still constrained by bundle sheath cell volume.

**Figure 7.**
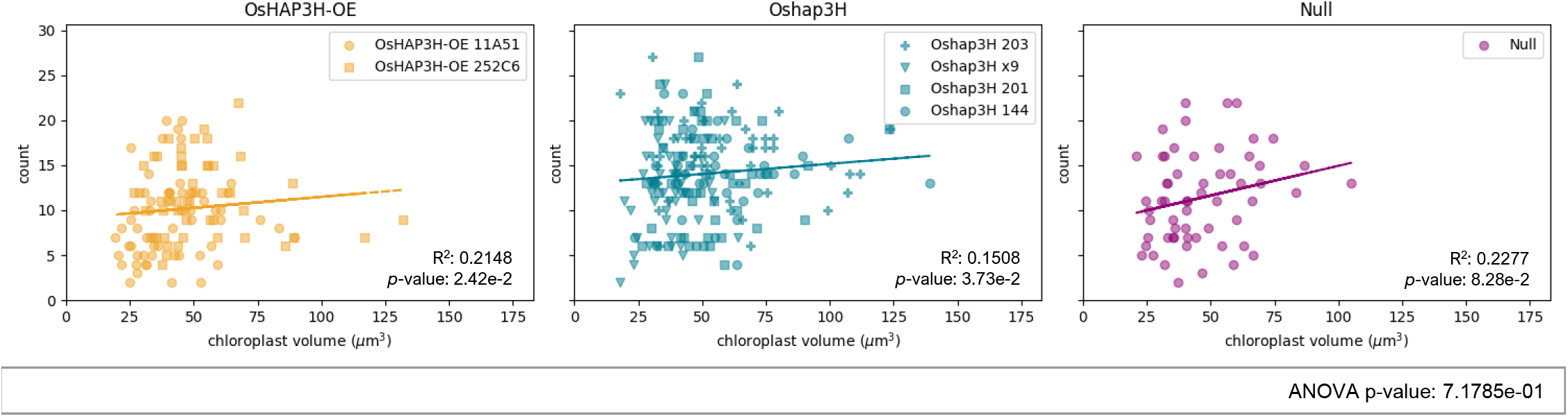
Correlation between chloroplast number and average chloroplast volume in *OsHAP3H-OE* and *Oshap3H* lines. Plots show untransformed data with corresponding linear regression depicted in solid line lines. R^2^ and p-values show variation and significance of the model for each analysis, respectively. Statistical analysis was based on transformed data to meet the parameters or parametric distribution. Each dot represents data from a single bundle sheath cell. An average of 10 cells from a minimum of four plants was assessed for each line. Lines with the same genetic background were grouped together and treated as one statistical group. Statistical comparison between groups was performed via ANOVA and p-value is shown at the bottom right. A p-value ≤ 0.05 would result in rejection of the null hypothesis that the groups are similar to each other. Details of statistical analyses are available in Table S1).

## DISCUSSION

### OsHAP3H represses chloroplast development in rice

The HAP/NF-Y/CBF proteins function as heterotrimeric complexes that regulate gene expression, with both HAP3/NF-YB and HAP5/NF-YC family members implicated in light-mediated developmental processes (Petroni *et al*., 2012). HAP2/NF-YA/CBF-B binds specifically to CCAAT boxes, whereas HAP3/NF-YB/CBF-A and HAP5/NF-YC/CBF-C have histone folding domains comparable to the core histones H2B and H2A, respectively (Petroni *et al*., 2012). Members of the *HAP5/NF-YC/CBF-C* group have been noted for their involvement in light-mediated developmental processes across various species. For instance, in Arabidopsis, *NF-YC* genes act synergistically with *HY5,* a key regulator of photomorphogenesis, to inhibit light-induced hypocotyl elongation (Myers *et al*., 2016). Similarly, in wheat, a significant proportion of NF-YC genes respond to light, and genes co-regulated with *TaNF-YC11* that have a CCAAT motif in their promoter region also influence chloroplast function (Stephenson *et al*., 2010). Given the coordinated action of NF-YC and NF-YB, it is unsurprising that most of the *NF-YB* members in wheat are also light-responsive (Stephenson *et al*., 2011). Overexpression of *TaNF-YB3* in wheat leads to increased chlorophyll accumulation, higher photosynthetic rates, taller stature and greater biomass (Stephenson *et al*., 2011). Despite both *OsHAP3H* and *TaNF-YB3* belonging to the *HAP3/NF-YB/CBF-A* group, overexpression of *OsHAP3H* in rice does not recapitulate the phenotype observed in wheat whereas loss of function enhances chlorophyll accumulation, tiller and panicles numbers and photosynthetic efficiency (quantified via ETR and LJPSII). OsHAP3H thus represses chloroplast development in rice whereas its counterpart in wheat promotes development.

The unexpected suppressor role for OsHAP3H in rice can be explained by the diverse mechanisms of action of *HAP/NF-Y/CBF* genes possibly acquired as a consequence of multiple gene duplication events within species that could lead to alteration of gene function. Specifically, HAP proteins can form heterotrimeric complexes with various compositions, including non-NF-Y members. For instance, the flowering regulator CONSTANS (CO) competes with NF-YA for the NF-YB/NF-YC dimer, increasing its affinity for DNA sequences other than the CCAAT motif (Gnesutta *et al*., 2017). Moreover, the NF-Y complex can act either as an activator or repressor of target gene expression. In cucumber, NF-YC2 and NF-YC9 upregulate the expression of *TIC21*, facilitating proper TOC-TIC complex function for protein import from the cytosol to the chloroplast (Ke *et al*., 2023). Conversely, in *Arabidopsis,* NF-YC proteins can interact with histone deacetylases (HDAs), which compact chromatin and repress gene expression (Tang *et al*., 2017). The precise mechanisms underlying the action of *OsHAP3H* in the bundle sheath cells of rice, or its effects in mesophyll cells, remains elusive. Nevertheless, it is evident that lack of *OsHAP3H* function activated chloroplast development in bundle sheath cells.

Analysis of tobacco and Arabidopsis mutants revealed that smaller and more numerous chloroplasts in mesophyll cells have a more positive impact on plant photosynthesis than fewer larger ones (Xiong *et al*., 2017; Głowacka *et al*., 2023). Smaller chloroplasts have relatively larger surface areas which enhances CO_2_ conductance, and they can move faster intracellularly to optimise light reception under fluctuating light conditions (Königer *et al*., 2008; Weise *et al*., 2015; Głowacka *et al*., 2023). Conversely, overexpression of maize *GOLDEN2* (*ZmG2)* in rice resulted in larger chloroplasts in bundle sheath cells, leading to higher carbon assimilation, biomass accumulation and seed yield in field-grown plants (Wang *et al*., 2013; Li *et al*., 2020). The evidence that loss of *OsHAP3H* function increases chloroplast numbers in bundle sheath cells of rice suggests that it acts as a repressor of chloroplast biogenesis. Recent work by Plackett and Hibberd (2024) supports the hypothesis that chloroplast biogenesis is normally repressed in rice bundle sheath cells. Similarly, we demonstrate that whereas the average chloroplast volume increases as bundle sheath cell dimensions increase, chloroplast numbers are constrained and thus chloroplast occupancy is reduced as cells get bigger. *OsHAP3H* has emerged as a candidate to increase chloroplast numbers in bundle sheath cells of rice, although the number constraining mechanism is still evident in larger cells of mutant plants. Future efforts to increase photosynthetic capacity in rice bundle sheath cells must overcome this constraint.

### Chloro-Count is a valuable tool for rapid quantification of chloroplast and bundle sheath dimensions in rice

The need for a tool enabling high-throughput characterization of chloroplast dimensions in rice bundle sheath cells led to the development of Chloro-Count. Previous comparisons of chloroplast volume quantification using images obtained through CLSM or SBF-SEM showed contrasting results. While some indicate that SBF-SEM is more reliable than CLSM (Lee *et al*., 2023), others support the robustness of CLSM imaging (Knoblauch *et al*., 2024). Chloro-Count utilizes deep learning to segment chloroplasts, identifying their occurrence and dimensions across z-stack sessions obtained via CLSM of isolated bundle sheath cells. Assessment of Chloro-Count’s efficiency to accurately segment chloroplasts and bundle sheath cells revealed high accuracy, particularly for bundle sheath cells, with segmentation producing 11% of false negatives and only 7.3% of false positives. These values were higher for chloroplasts, at 38% and 23%, respectively. Despite these relatively low figures, an additional step of manual verification of segmentation was introduced to further enhance the accuracy of chloroplast and bundle sheath cell segmentation. With this step, Chloro-Count proved to be a valuable tool for precise identification of chloroplasts and bundle sheath cells in CLSM images in rice.

We have demonstrated that accurately estimating chloroplast and bundle sheath cell dimensions in 2D sections is challenging, likely due to the various shapes and differential distribution along the cell axis. Similarly, chloroplast numbers in mesophyll cells of Norway spruce needles were 10 times underestimated in 2D images compared to 3D images (Kubínová *et al*., 2014). Therefore, quantification considering all cell dimensions provides a more accurate calculation of chloroplast occupancy of bundle sheath cells. Previously, volume estimation based on 2D sections was conducted for wheat and chickpea mesophyll and bundle sheath cells (Harwood *et al*., 2020). Chloroplast volume was calculated using the ellipsoid volume formula, based on the height and length of chloroplasts of both cell types (Harwood *et al*., 2020). However, compared to 3D reconstructions of the cell via SBF-SEM, both cell and chloroplast volumes were underestimated when solely based on 2D sections (Harwood *et al*., 2020). In the present study, volumetric estimation based on ellipsoid reconstruction through z-stacks showed significant variation and the approach was discarded. Instead, Chloro-Count quantifies volumes based on the sum of area multiplied by the distance between z-stacks, a simple approach that accounts for the variability in chloroplast shapes and the position of chloroplasts within the bundle sheath cell.

Previous studies have quantified chloroplasts using methods based on machine learning. DeepLearnMOR was a model trained to differentiate the morphology of normal and aberrant chloroplast in *arc6-5* mutants in Arabidopsis (Li *et al*., 2021). Conversely, Feng et al. (2023) reconstructed chloroplast 3D structures based on SBF-TEM sections, while Su et al. (2023) described a pipeline to quantify the number of chloroplasts in bright field images from three species of bryophytes. However, none of these studies combine the quantification of both the volume and number of chloroplasts in a high throughput manner. In that context, Chloro-Count emerges as a valuable tool for rapid quantification of chloroplasts in rice bundle sheath cells, with precision comparable to similar methods using confocal imaging (Wolny *et al*., 2020; Khan *et al*., 2020; Vijayan *et al*., 2024). Its utilization with CLSM images, requiring minimal sample preparation, facilitates a high-throughput workflow. This allowed for accurate quantification of the increased chloroplast number in *Oshap3H* mutants, offering a novel route to augment chloroplast occupancy in bundle sheath cells alongside the previously characterised lines in which chloroplast size was enhanced. Collectively, this study provides valuable insights and resources for enhancing photosynthetic capacity in rice.

## MATERIALS AND METHODS

### Phylogenetic reconstruction

Rice OsHAP3 sequences were individually used as bait to identify orthologs using SHOOT (https://www.shoot.bio/; Emms and Kelly 2022). Sequences from *Zea mays*, *Oryza sativa, Triticum aestivum* and *Arabidopsis thaliana* were selected along with a single sequence from *Chlamydomonas reinhardtii* for further analysis. After manual removal of duplicates, sequences were aligned with MAFFT using the L-INS-i refinement method (Katoh et al. 2019, File S1). IQTree (Trifinopoulos *et al*., 2016) was used to estimate the best-fitting model parameters (JTT+F+I+G4) and to infer a consensus phylogenetic tree from 1000 bootstrap replicates. The data were imported into ITOL (Letunic & Bork, 2007) to generate the pictorial representation. The tree was rooted with the *C. reinhardtii* sequence and branches with less than 50% bootstrap support were deleted.

### Construct design and assembly

The coding sequence from *O. sativa HAP3H* (LOC_Os08g07740.1) was modified by adding silent mutations every ∼10 bp along the nucleotide sequence to avoid RNA-mediated transcriptional silencing and to facilitate the detection of transcripts from the transgene. It was further codon optimized for rice (cdo) and domesticated to remove any recognition sites for type II restriction enzymes used on Golden Gate cloning steps: BsaI, BpiI, Esp3I and DraIII. The final sequence was synthesised as a level 0 module with appropriate SC flanking sites (Engler & Marillonnet, 2013).

Cloning was carried out using standard Golden Gate parts and the one-step one-pot protocol (Engler *et al*., 2009). *OsHAP3H*_cdo_ module (EC17576) was cloned downstream of the maize ubiquitin promoter (*ZmUBI*_pro_, EC15455) and upstream of a nos terminator (tNOS, EC41421) on a level 1 vector position forward 2 (pICH47742). The previously described *hygromycin phosphotransferase* (HygR) coding sequence cloned downstream of the rice actin promoter (*OsACT*_pro_) was used as the selectable marker (Vlad *et al*., 2019). Level 1 modules were assembled into the binary vector pAGM4723 to obtain the construct EC17633 depicted in Figure S1.

### Plant Transformation

To obtain stable transgenic rice lines, the construct EC17633 was transformed into *Agrobacterium tumefaciens* strain EHA105 and co-cultivated with calli from seeds from *Oryza sativa spp. japonica* cultivar Kitaake. Callus transformation and seedling regeneration were performed according to a protocol modified from Toki et al. (2006), that can be downloaded at https://langdalelab.files.wordpress.com/2015/07/kitaake_transformation_2015.pdf.

Transformed T_0_ plantlets were verified by polymerase chain reaction (PCR) using primers to detect the presence of the selectable marker gene HygR (JLF_Ox12 and JLF_Ox13, Table S2). Positive plants were transferred to substrate and further analysed for T-DNA copy number.

### Plant growth

Rice seeds were de-hulled and sterilized with 80% ethanol for 2 minutes followed by 25% bleach for 25 minutes. Seeds were thoroughly washed with sterilized ddH2O before being sown on germination media (2.15 g/L Murashige and Skoog medium including vitamins – Duchefa-Biochemie, 44 mM sucrose, 2.5 mM MES, 2 g/L Phytagel, pH 5.8). Seeds were placed in an incubator with 16/8h photoperiod and temperature varying from 30°C during daytime to 24°C at night.

After two weeks, seedlings were transferred to pots with clay granules (Porous Ceramic Topdressing and Construction Material, Profile, UK) and grown in a controlled environment room with 16/8h light/dark regime, average light intensity of 300 μmol/m^2^/s, day temperature of 28°C and night of 25°C. Plants were irrigated with fertilizer solution (1.3 g/L Peters Excel Cal-Mag Grower N.P.K. 15-5- 15, Everris, UK) and supplemented with 7% chelated iron every two weeks until flowering.

### DNA blot analysis

Rice leaf tissue was ground in liquid nitrogen and genomic DNA (gDNA) was extracted using 500 μl of CTAB extraction buffer (1.5% CTAB, 75 mM Tris-HCl pH 8.0, 15 mM EDTA, 1.05 M NaCl). After incubation at 65°C for 20 minutes, samples were thoroughly mixed with an equal volume of Chloroform: Isoamylalcohol (24:1) and centrifuged at 13000 rpm for 10 minutes. The aqueous phase was mixed with the same volume of isopropanol for 10 minutes at room temperature for DNA precipitation. After centrifugation at the same conditions described above, the pellet was washed with 70% ethanol, air-dried and resuspended in 50 μl of ddH2O.

For each transgenic plant, 10 μg of gDNA was digested with SacI restriction endonuclease (New England Biolabs). Digested samples were electrophoresed on a 1% agarose gel stained with SYBR Safe (Invitrogen) for 16h at 20V. The agarose gel was washed with Denaturation solution (0.5M NaOH, 1.5M NaCl) for 30 minutes, followed by two washes with Neutralizing solution (0.5 M Tris pH 7.5, 1.5M NaCl) for 20 minutes each. Sheared gDNA was transferred overnight onto Hybond N+ membrane (GE Healthcare, UK) by capillary action.

After transfer, the membrane was UV crosslinked and hybridized for 16h with a digoxygenin (DIG)- labelled probe specific for the HygR gene (JLF_Ox41 and JLF_Ox42, Table S2) in DIG Easy Hyb solution (Merck). Subsequent steps included stringency washes to remove any spurious annealing or unbound probe. The membrane was blocked for 2h at room temperature with 3% skimmed milk w/v in 100 mM Maleic Acid, 150 mM NaCl (pH 7.5), followed by 30 minutes incubation with Anti-Digoxigenin-AP, Fab fragments (Merck). Signals were detected using CDP-Star according to the manufacturer’s instructions (Roche Diagnostics).

### Gene expression analysis via qPCR

The mid-portion of leaf 4 from 20 days after sowing rice plants was harvested and immediately frozen in liquid nitrogen. Ground tissue was used for RNA extraction using the RNeasy Plant Mini Kit (Qiagen) according to manufacturer’s instructions. To avoid any DNA contamination, samples were treated with TURBO DNA-free DNase (Thermo Fischer Scientific) according to the manufacturer’s guidelines. For first strand cDNA synthesis, 300 ng of total RNA was added to 1 μl of Maxima Enzyme Mix (Thermo Fisher Scientific) and 1x Reaction Mix to a final volume of 10 μl. The mixture was incubated at 25°C for 10 minutes, followed by 50°C for 15 minutes. Enzyme inactivation was induced by incubation at 85°C for 5 minutes.

The amplification reactions were performed in a StepOnePlus Real-Time PCR System (Thermo Fisher Scientific) using SYBRGreen to monitor dsDNA synthesis. The reaction mixtures contained 5 μl of diluted cDNA (1:25) and 10 μl of SYBR™ Green PCR Master Mix (Thermo Fisher Scientific) in a total volume of 20 μl. The reaction cycles began with a five-minute denaturation step at 94°C, followed by 40 amplification cycles of 15 seconds each at 94°C, 10 seconds at 60°C, 15 seconds at 72°C. After each cycle, the fluorescence was measured at 60°C for 35 seconds. The melting curve was produced by a cycle of 95°C for 15 seconds, 60°C for one minute, 95°C for 30 seconds and finally 60°C for 15 seconds. Each single qPCR reaction was repeated three times to make technical replicates.

The expression level of *OsHAP3H*_cdo_ and *OsHAP3H_e_*was assessed with primers designed to specifically amplify each allele (JLF_Ox131 and JLF_Ox132, and JLF_Ox176 and JLF_Ox177, respectively; Table S2). Two reference genes were used to normalize gene expression levels, *OsACT* (Actin-F and Actin-R, Wang et al., 2017, Table S2) and *OsUBQ5* (JLF_Ox161 and JLF_Ox162, Jain et al., 2006, Table S2). Exported *Rn* values were used to calculate amplification efficiency for each primer pair and the Cq values for each qPCR reaction using the ‘*qpcR*’ package from R (Ritz & Spiess, 2008). Normalized relative expression of the target genes was calculated using the R package ‘*EasyqpcR’* (https://www.bioconductor.org/packages//2.12/bioc/html/EasyqpcR.html), according to the ΔΔCq model (Pfaffl, 2001) based on the previously calculated Cq and amplification efficiencies.

### Total chlorophyll extraction and quantification

Three leaf discs were harvested from leaf 7 of plants from 33 DAS and immediately frozen in liquid nitrogen. For total chlorophyll extraction, finely powdered tissue was resuspended in 80% acetone buffered in 100 mM Tris-HCl pH 8.0 (Chazaux *et al*., 2022). Following a centrifugation step, the absorbance of each sample was read in technical triplicates in a FLUOstar Omega spectrometer (BMG Labtech) at 646 and 663 nm. Background absorbance was measured at 750 nm. Absorbance values recorded at 750 nm were subtracted from values recorded at 646 and 663 nm for normalization purposes. Technical replicates were averaged for each sample. Concentration of chlorophylls a and b was calculated in micrograms (μg) per leaf area using the following the equations (13.71 x Abs_663_ - 2.85 x Abs_646_) and (22.39 x Abs_646_ - 5.42 x Abs_663_), respectively (Porra *et al*., 1989).

### Field Trial

Seeds from *Oshap3H* lines alongside null and Kitaake controls were germinated in the laboratory and after 12 days, 24 plants per line were transferred to randomized plots in the field at the GM Experiment Station in NCHU, Wufeng District, Taichung City, Taiwan during the period of August to November of 2022. After 50 days growing in the field, 12 plants per line were analysed for chlorophyll fluorescence parameters. Total chlorophyll was measured using a SPAD 502 Plus Chlorophyll Meter (Spectrum Technologies Inc.). The local radiance (PAR) and the photosynthetic parameters ETR and LJPSII were measured with a MINI-PAM Portable Chlorophyll Fluorometer (Heinz Walz GmbH). At the end of the season, measurements of plant height, number of tillers, above ground biomass excluding panicle and panicle weight, number and length were taken. Seed weight was measured by weighing 1000 seeds. Total biomass was calculated by combining above ground biomass and panicle weight. Yield was estimated by measuring grams of seeds generated per plant.

### Bundle sheath cell image acquisition

The mid-portion of fully expanded leaf 7 from 43 days after sowing (DAS) rice plants were sectioned longitudinally in between two lateral veins and fixed in 0.5% glutaraldehyde in phosphate saline buffer pH 7.4 for 16h at 4°C. Samples were then incubated in 0.2M Na-EDTA pH 9.0 at 55°C for 3h. Leaf digestion was performed by incubating samples in *Aspergillus niger* pectinase (Sigma-Aldrich, cat no. P4716) at a 2.5% v/v dilution in digestion buffer (0.15M H2NaPO4, 0.04M Citric Acid, pH 5.3) for 2h at 45°C. Samples were then rinsed twice in digestion buffer and stored at 4°C until imaging.

Upon imaging, leaf samples were physically loosened in a glass slide to release bundle sheath strands and mounted in 50% glycerol under a coverslip. Images of bundle sheath cells were taken with 20x magnification with a water immersion objective lens on a Leica SP5 confocal microscope. Digital zoom was used to centralise one bundle sheath cell per image. Chlorophyll autofluorescence was obtained at 633 nm excitation and 650-750 nm emission wavelengths with gain of 767V and offset of 0.1%. Bundle sheath cell outlines were obtained in the bright field channel with gain ranging from 265-283V and offset of −4.4%. Acquired images were 512x512 pixels in size and two consecutive z-stacks spanned 0.99 μm. All images were exported from the microscope as .lif files with metadata that included pixel size of each image.

### Chloro-Count Overview

Chloro-Count is used to identify and measure volumes for bundle sheath cells and individual chloroplasts. This process follows a semi-supervised approach, where two independently trained image segmentation networks detect contours for organelles and cell boundaries in each cell slice. The detected regions are represented as a list of connected polygons, facilitating visual correction and validation from an expert. Once individual segments have been validated, they are mapped to individual organelles/cells for volumetric analysis. An overview of the system for detecting and measuring volumes of chloroplasts is presented in Figure 3A. The process for detecting and measuring bundle sheaths follows an analogous workflow. The Chloro-Count code is available on https://github.com/pedropgusmao/chloro-count.

### Data collection and pre-processing

A total of 327 slices from 39 different cells were used during the training of both image segmentation networks. Images from 29 cells were used for training, five for validation and five for testing. A total of 3,790 segments of chloroplasts were used for training, 287 for validation and 599 for testing. Bundle sheath cells used 327 segments for training (one per slice), 65 for validation and 60 for testing. Images were manually annotated by two experts for bundle sheath cells and chloroplasts using VGG Image Annotator v. 1.0.6 (https://www.robots.ox.ac.uk/~vgg/software/via/via-1.0.6.html). Unique labels were created for individual chloroplasts to track them along the z-stacks within a cell, allowing us to validate our aggregation strategy during volumetric analysis.

### Image Segmentation and Segment Association

We used two Mask R-CNN models with a ResNet50 Feature Pyramid Network backbone from Torchvision (https://github.com/pytorch/vision) to produce region candidates for both bundle sheath and chloroplast targets. The networks’ last layers were modified to include a binary classifier and both networks were trained for ten epochs using a Stochastic Gradient Decent optimiser with learning rate of 5e-3, momentum 0.9, and weight decay of 5e-4. A learning rate scheduler was also used with step size 3 and gamma 0.1 to improve training.

Our fully-trained models output candidate regions and confidence scores for each suggested segmentation. These proposed regions often overlap and are filtered using non-maxima suppression. The network responsible for segmenting bundle sheaths produced an AR of 0.768 on the holdout test set considering an Intersection over Unions (IoU) of 0.5-0.95 and an AP of 0.927 (@IoU=0.75). The network responsible for segmenting chloroplasts produced an AR of 0.623 (@IoU=0.5-0.95) and AP of 0.89 (@IoU=0.75) on the same set and for the same range of IoU. We emphasise the use of Average Recall (AR) as our main metric, as it is easier for annotators to delete false positives than to create new annotations during the verification stage. The code for segmenting chloroplasts and bundle sheath cells is available on https://github.com/pedropgusmao/chloro-count.

### Volume and Area Estimations

After the segments detected by the neural networks have been validated or adjusted, the area of each slice can be directly inferred from their internal number of pixels. However, to measure volumes, we still need to associate each segment with individual chloroplasts. In the case of bundle sheath cells, this association is trivial as there is only one segmented bundle sheath per cell slice. Aggregation of contours as individual chloroplast is done based on the distance between centres of mass of contours across image slices. The plane containing the highest number of chloroplasts is used as an initial reference. Segments found on planes immediately above and below the reference plane are matched to distinct segments whose X-Y centre of mass coordinates are closest. An empirical threshold is set to prevent matches between segments which are far apart.

Once all segments had been associated with specific chloroplasts, we proceeded with volumetric evaluation. We explored three different methods for estimating the volume of both bundle sheath cells and chloroplasts. The simplest approach for volume quantification was to calculate, for each chloroplast, the sum of the areas of each segment times the distance between cell slices. The code for counting and measuring chloroplasts and bundle sheath cells is available on https://github.com/pedropgusmao/chloro-count.

### Statistical analysis

For comparisons between 2-D and 3-D measurements and the effects of bundle sheath cell in chloroplasts dimensions, each parameter analysed had its normal distribution confirmed via a Shapiro-Wilk test (Shapiro & Wilk, 1965). Those parameters that did not follow a normal distribution were transformed by Log10 (Table S1). Linear regressions were performed using the library SciPy v.1.11.1 to verify how different parameters correlated to each other, and R^2^ and p-values were used to assess the variation and the significance of the model, respectively. A p-value lower than 0.05 was the threshold to reject the null hypothesis that the linear regression slope equals zero (the dependent variable is constant and therefore, not dependent on the independent variable). Both R^2^ and p-values calculated based on parametric data are depicted on each figure.

For relative gene expression, statistical significance was calculated based on a Student’s t-test, comparing the mean variance of each mutant line with the null (Table S1). For bundle sheath cell and chloroplast volumes, chloroplast number, relative occupancy and chlorophyll concentration, the Shapiro-Wilk test was used to evaluate normality, and for those deviating from that either square root or boxcox transformations were applied (Table S1). The difference between group means was calculated via a two-way ANOVA followed by a pairwise comparison analysis with Tukey’s HSD test (Tukey, 1949) (Table S1). A p-value lower than 0.05 was the threshold to reject the null hypothesis that the means between groups did not differ. The Student’s t-test was performed using the library SciPy v.1.11.1, while the ANOVA and Tukey’s HSD analysis were performed using statsmodels v.0.14.0 libraries. Plots were prepared using matplotlib v.3.7.1 library from Python v.3.11.6; using Visual Studio Code as code editor.

## AUTHOR’S CONTRIBUTION

JL-F and JAL conceived and designed the experiments; RWH and SK contributed to initial experimental design; PPBG designed and created the Chloro-Count model, and performed the quantification of chloroplasts and bundle sheath cells; GS carried out plant transformation, cultivation and genotyping; S-FL and S-MY performed and collected the data from the field experiment; RWH and SK generated the silent mutations along *OsHAP3H_cdo_* coding sequence. JL-F carried out all of the remaining experiments and analysed the data. JL-F, PPBG and JAL wrote the manuscript with edits of the penultimate version contributed by RWH and SK.

## ACKNOWLEDGEMENTS

The Authors thank Roxaana Clayton, Julie Bull and Lizzie Jamison for technical support; Anna Hermanns for assisting with initial characterization of *OsHAP3H*-OE lines; Andy Plackett for providing guidance on the bundle sheath isolation protocol and comments on statistical analysis; Sophie Johnson, Chiara Perico, Daniela Vlad, Sovanna Tan, Thomas Hughes, Maricris Zaidem and Julian Hibberd for discussion throughout the experimental work.

---

**Figure S1.** Genotype of *OsHAP3H-*OE and *Oshap3H* lines. **A)** Schematic of the construct used to over-express *OsHAP3H_cdo_. HygR* depicts the hygromycin phosphotransferase gene and *OsACT_pro_*and *ZmUBI_pro_* represent the constitutive rice actin and maize ubiquitin promoters, respectively. LB and RB refer to left and right borders. The position of the hybridization probe used for DNA blot analysis is depicted as a black bar. The parallel arrows indicate the amplicon site used for qPCR amplification. **B)** Alignment between the native *OsHAP3H_n_* and optimized *OsHAP3H_cdo_* nucleotide sequences. Light blue shading highlights the sites of silent mutations; the red square indicates the site of CRISPR gRNA (gRNA 42) targeting; the solid and dashed arrows show the primer binding sites for qPCR amplification of *OsHAP3H_e_* and *OsHAP3H_cdo_,* respectively. **C)** DNA gel blot analysis of null segregant and *OsHAP3H-OE* lines digested with the SacI restriction enzyme which cuts at two sites between the hybridization probe and the right border (see S1A), and hybridized to a fragment of the HygR gene. Two independent transgene insertions are evident in each *OsHAP3H-OE* line. **D)** Wild-type (WT) and mutant sequences with the predicted protein length. Each line represents a single transformation event.

**Figure S2.** Relationship between two-dimensional cell parameters and cell volume. **A)** Schematic showing how different bounding box shapes (representing a bundle sheath cell) can have the same face area (representing a z-stack) yet very different volumes. Differences in x/y ratios, however, are similar to volume differences. **B)** Correlation between cell volume and bounding box x/y ratio. Grey lines show linear regression of data obeying a normal distribution; R^2^ and p-values show variation and significance of the model for each analysis, respectively. Details of statistical analyses are available in Table S1).

**Figure S3.** Distribution of chloroplasts along 3-dimensional cell axes. **A)** Transverse section of a rice leaf showing an intermediate vein (v) surrounded by a ring of bundle sheath cells (bs). Magenta colour is chlorophyll autofluorescence in chloroplasts, and green is propidium iodide staining of the cell wall. Scale bar = 50 µm. **B)** Schematic figure of cell axes. A bundle sheath cell is represented by a blue cylinder cut by two planes. Where the planes cross is the centre of the cell (zero). Cell length in the proximo-distal leaf axis is represented by ‘x’, cell height in the adaxial-abaxial leaf axis is represented by ‘y’ and cell width in the medio-lateral leaf axis is represented by ‘z’. Images for quantification were taken sequentially through the z axis with x as the face. **C-E)** Distribution of chloroplasts along the x (B), y (C) and z (D) axes of the cell depicted as a histogram with 1500, 1500 and 75 bins, respectively. The vertical axis of the plot indicates the probability density, which is the count of chloroplasts in that bin divided by the total number of counts in all bins. The horizontal axis of the plot indicates the relative distance from the centre of the bundle sheath cell (zero). The red line shows a Gaussian kernel density estimate (KDE), a density estimation derived from the data that models the distribution of points.

**Table S1.** Statistical analysis.

**Table S2.** PCR Primer Sequences.

File S1. Protein sequences of HAP gene family used for phylogenetic reconstruction.

