## Supplementary material for "INCREASED CHLOROPLAST OCCUPANCY IN BUNDLE SHEATH CELLS OF RICE *hap3H* MUTANTS REVEALED BY CHLORO-COUNT, A NEW DEEP LEARNING-BASED TOOL": Figure S1

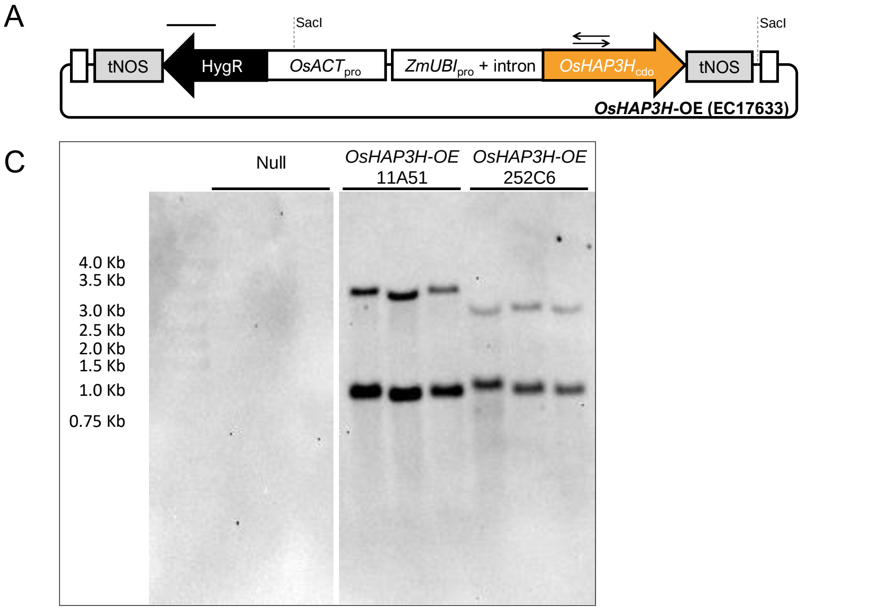


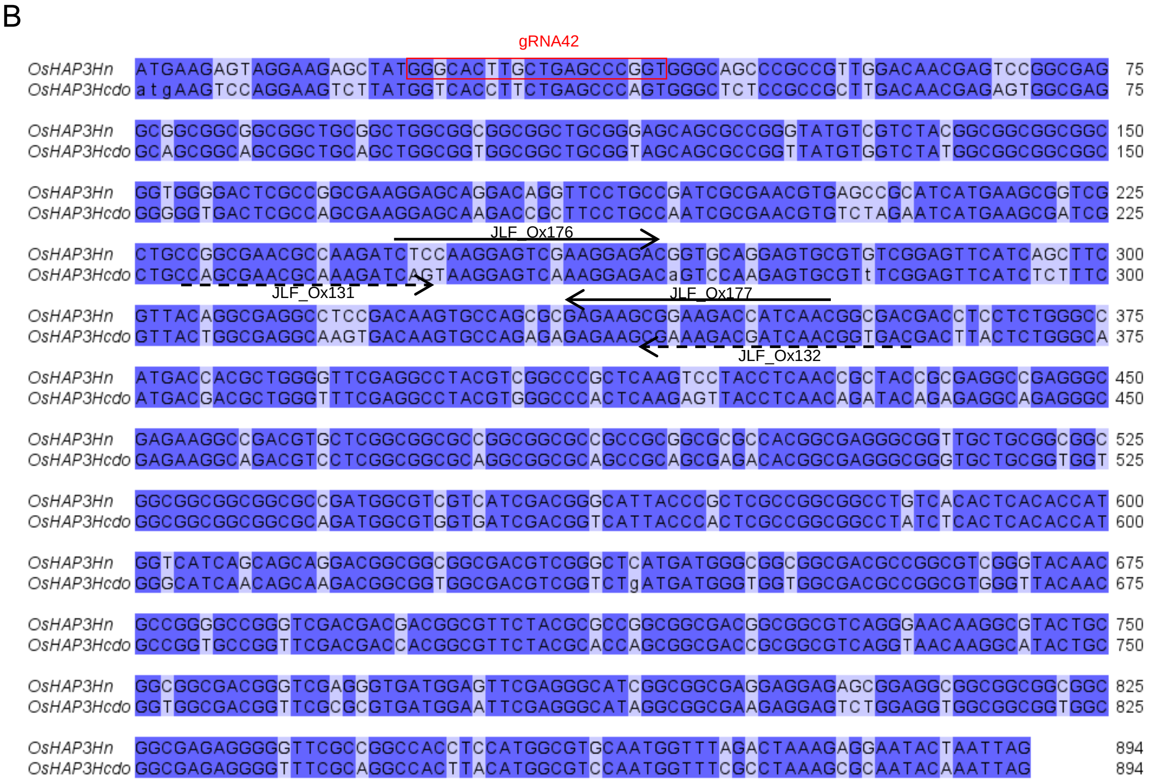


D

| Line | Sequence | Putative protein length |
|---|---|---|
| WT | ATGAAGAGTAGGAAGAGCTATGGGCACTTGCTGAGCCC-GGTGGGCAGCCCGCCGTTGGA | 297 aa |
| *Oshap3H* 144 | ATGAAGAGTAGGAAGAGCTATGGGCACTTGCTGAGCCCTGGTGGGCAGCCCGCCGTTGGA | 290 aa |
| *Oshap3H* 201 | ATGAAGAGTAGGAAGAGCTATGGGCACTTGCTGA--------GGGCAGCCCGCCGTTGGA | 65 aa |
| *Oshap3H* 203 | ATGAAGAGTAGGAAGAGCTATGGGCACTTGCTGAGC---GGTGGGCAGCCCGCCGTTGGA | 289 aa |
| *Oshap3H* x9 | ATGAAGAGTAGGAAGAGCTATGGGCACTTGCTGAGCC--GGTGGGCAGCCCGCCGTTGGA | 67 aa |

**Figure S1. Genotype of *OsHAP3H*-OE and *Oshap3H* lines. A)** Schematic of the construct used to over-express $OsHAP3H_{cdo}$. *HygR* depicts the hygromycin phosphotransferase gene and $OsACT_{pro}$ and $ZmUBI_{pro}$ represent the constitutive rice actin and maize ubiquitin promoters, respectively. LB and RB refer to left and right borders. The position of the hybridization probe used for DNA blot analysis is depicted as a black bar. The parallel arrows indicate the amplicon site used for qPCR amplification. **B)** Alignment between the native $OsHAP3H_n$ and optimized $OsHAP3H_{cdo}$ nucleotide sequences. Light blue shading highlights the sites of silent mutations; the red square indicates the site of CRISPR gRNA (gRNA 42) targeting; the solid and dashed arrows show the primer binding sites for qPCR amplification of $OsHAP3H_e$ and $OsHAP3H_{cdo}$, respectively. **C)** DNA gel blot analysis of null segregant and *OsHAP3H-OE* lines digested with the SacI restriction enzyme which cuts at two sites between the hybridization probe and the right border (see S1A), and hybridized to a fragment of the HygR gene. Two independent transgene insertions are evident in each *OsHAP3H-OE* line. **D)** Wild-type (WT) and mutant sequences with the predicted protein length. Each line represents a single transformation event.
