## Supplementary material for "INCREASED CHLOROPLAST OCCUPANCY IN BUNDLE SHEATH CELLS OF RICE *hap3H* MUTANTS REVEALED BY CHLORO-COUNT, A NEW DEEP LEARNING-BASED TOOL": Figure S2

A

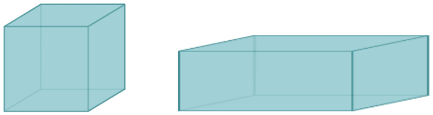

| x axis | 4 | 8 |
|---|---|---|
| y = z axis | 4 | 2 |
| Face area | 16 | 16 |
| Volume | 64 | 32 |
| x/y ratio | 1 | 2 |

B

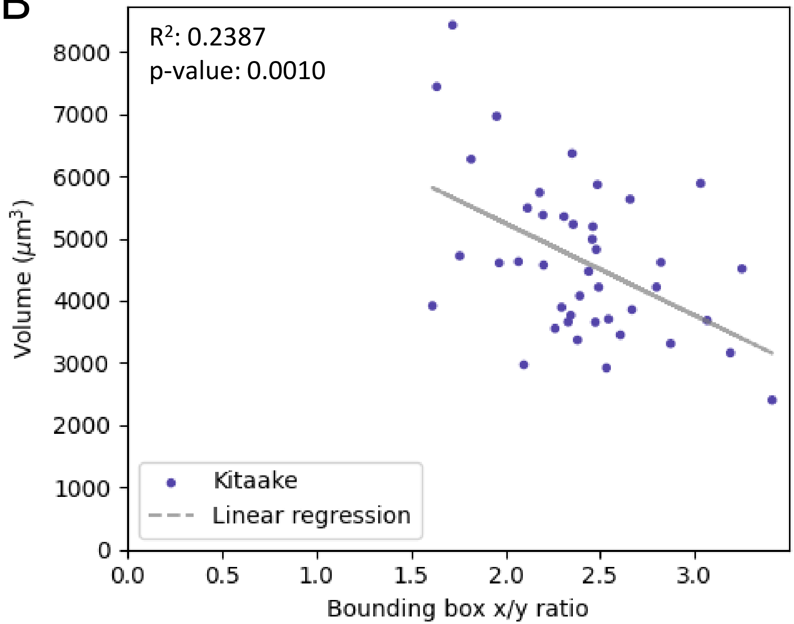


**Figure S2. Relationship between two-dimensional cell parameters and cell volume. A)** Schematic showing how different bounding box shapes (representing a bundle sheath cell) can have the same face area (representing a z-stack) yet very different volumes. Differences in x/y ratios, however, are similar to volume differences. **B)** Correlation between cell volume and bounding box x/y ratio. Grey lines show linear regression of data obeying a normal distribution; $R^2$ and p-values show variation and significance of the model for each analysis, respectively. Details of statistical analyses are available in Table S1).
