## Supplementary material for "INCREASED CHLOROPLAST OCCUPANCY IN BUNDLE SHEATH CELLS OF RICE *hap3H* MUTANTS REVEALED BY CHLORO-COUNT, A NEW DEEP LEARNING-BASED TOOL": Figure S3

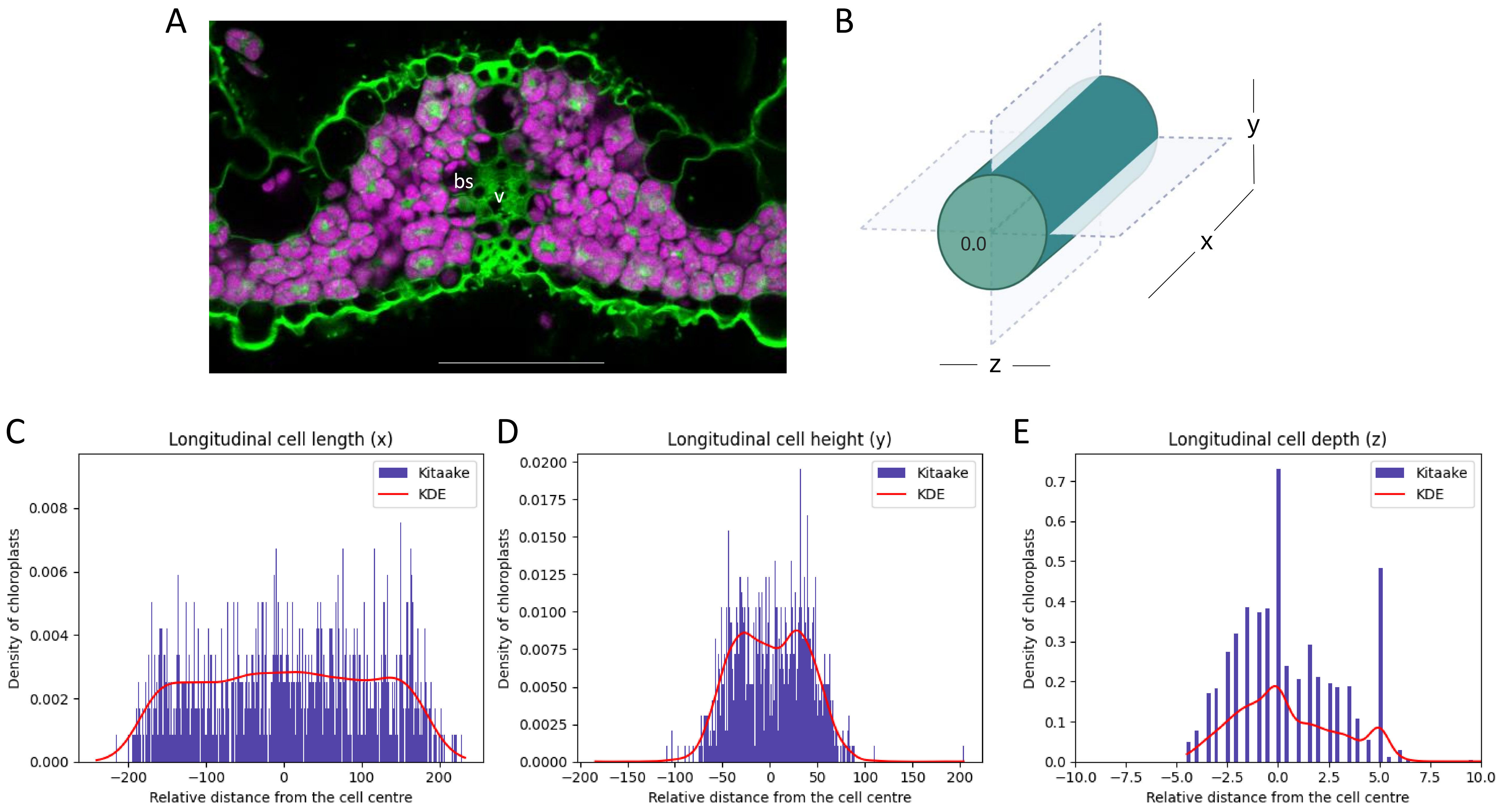


**Figure S3. Distribution of chloroplasts along 3-dimensional cell axes. A)** Transverse section of a rice leaf showing an intermediate vein (v) surrounded by a ring of bundle sheath cells (bs). Magenta colour is chlorophyll autofluorescence in chloroplasts, and green is propidium iodide staining of the cell wall. Scale bar = 50 µm. **B)** Schematic figure of cell axes. A bundle sheath cell is represented by a blue cylinder cut by two planes. Where the planes cross is the centre of the cell (zero). Cell length in the proximo-distal leaf axis is represented by ‘x’, cell height in the adaxial-abaxial leaf axis is represented by ‘y’ and cell width in the medio-lateral leaf axis is represented by ‘z’. Images for quantification were taken sequentially through the z axis with x as the face. **C-E)** Distribution of chloroplasts along the x (B), y (C) and z (D) axes of the cell depicted as a histogram with 1500, 1500 and 75 bins, respectively. The vertical axis of the plot indicates the probability density, which is the count of chloroplasts in that bin divided by the total number of counts in all bins. The horizontal axis of the plot indicates the relative distance from the centre of the bundle sheath cell (zero). The red line shows a Gaussian kernel density estimate (KDE), a density estimation derived from the data that models the distribution of points.
