## Supplementary material for "INCREASED CHLOROPLAST OCCUPANCY IN BUNDLE SHEATH CELLS OF RICE *hap3H* MUTANTS REVEALED BY CHLORO-COUNT, A NEW DEEP LEARNING-BASED TOOL": File S1

**Sequences**

>OsHAP3A_LOC_Os01g61810.2

MMMMDLGFLEGGAGMADAGHDESGSPPRSGGVREQDRFLPIANISRIMKKAVPANGKIAK

DAKETLQECVSEFISFVTSEASDKCQKEKRKTINGEDLLFAMGTLGFEEYVDPLKIYLHK

YRELVG*

>OsHAP3J_LOC_Os01g70880.1

MTNGQDNLLPIANVGRIMKDGLPPQAKISKRAKETIQECATEFISFVTGEASERCRRERR

KTVNGDDVCHAMRSLGLDHYADAMHRYLQRYREGEELAASLNSSSSAAAAAAAAGSRGGG

AIQIDVRAELSIFRSGNNQGRPNN*

>OsHAP3G_LOC2_Os01g70890.1

MADHHGGHHADGHRRQQQLQGEAADQAAAEIIKEQDRLLPIANVGRIMKQILPPNAKISK

EAKETMQECVSEFISFVTGEASDKCHKEKRKTVNGDDVCWAFGALGFDDYVDPMRRYLNK

YRELEGDRAAAAATSRSGAGAAAGPDHPSSSSSAAAATAGHFMFNAMDRSTDSSRQF*

>OsHAP3E_LOC_Os02g49370.1

MEAGYPGTAANGAAADGNGGAQQAAAAPAIREQDRLMPIANVIRIMRRVLPAHAKISDDA

KETIQECVSEYISFITGEANERCQREQRKTITAEDVLWAMSRLGFDDYVEPLGVYLHRYR

EFEGESRGVGVGVGAARGDHHHGHVGGMLKSRAQGSMVTHHDMQMHAAMYGGGAVPPPPH

PPPHHHAFHQLMPPHHGQYAPPYDMYGGEHGMAAYYGGMYAPGSGGDGSGSSGSGGAGTP

QTVNFEHQHPFGYK*

>OsHAP3K_LOC2_Os02g49410.1

MAGNKKRGGRNMDQVKKAAVRSDGVGGSATNAELPMANLVRLIKKVLPGKAKIGGAAKGL

THDCAVEFVGFVGDEASEKAKAEHRRTVAPEDYLGSFGDLGFDRYVDPMDAYIHGYREFE

RAGGNRRVAPPPPAAATPLTPGGPTFTDAELQFLRSVIPSRSDDEYSGSSPAIGGYGYGY

GYGKNM*

>OsHAP3I_LOC_Os03g29970.1_

MPDSDNDSGGPSNYAGGELSSPREQDRFLPIANVSRIMKKALPANAKISKDAKETVQECV

SEFISFITGEASDKCQREKRKTINGDDLLWAMTTLGFEDYVDPLKHYLHKFREIEGERAA

ASTTGAGTSAASTTPPQQQHTANAAGGYAGYAAPGAGPGGMMMMMGQPMYGSPPPPPQQQ

QQQHHHMAMGGRGGFGHHPGGGGGGSSSSSGHDEDEDEDEDEDED*

>OsHAP3B_LOC_Os05g38820.4

MADGPGSPGGGGGSHESGSPRGGGGGGGGGGGGGGVREQDRFLPIANISRIMKKAIPANG

KIAKDAKETVQECVSEFISFITSEASDKCQREKRKTINGDDLLWAMATLGFEDYIEPLKV

YLQKYREVRTDVDVWNWGDSLLI*

>OsHAP3C_LOC_Os05g49780.1

MSEGFDGTENGGGGGGGGVGKEQDRFLPIANIGRIMRRAVPENGKIAKDSKESVQECVSE

FISFITSEASDKCLKEKRKTINGDDLIWSMGTLGFEDYVEPLKLYLRLYREGDTKGSRAS

ELPVKKDVVLNGDPGSSFEGM*

>OsHAP3D_LOC_Os06g17480.1

MEPAFPNGGAAAPPPPMAAEQLPPAAAVVREQDRLMPIANVIRIMRRVLPPHAKISDDAK

EVIQECVSEFISFVTGEANDRCHREHRKTVTAEDLVWAMDRLGFDDYVPPLTAYLRRMRE

YEGGGSGGGGGGGRGAAAAPAVVPPPPPPPPEDAFRYVQVHHPVYAAPGEPVQGYGYPVA

MSSALPAPHVHVGVRGGGQHEVFGGGPAPLAVYYGGAPYGEASSRGGCSAADEGSSSSSA

SPAPVGPNYE*

>OsHAP3F_LOC_Os07g41580.1

MPDSDNESGGPSNAGEYASAREQDRFLPIANVSRIMKRALPANAKISKDAKETVQECVSE

FISFITGEASDKCQREKRKTINGDDLLWAMTTLGFEDYIDPLKLYLHKFRELEGEKAIGA

AGSGGGGAASSGGSGSGSGSHHHQDASRNNGGYGMYGGGGGMIMMMGQPMYGSPPASSAG

YAQPPPPHHHHHQMVMGGKGAYGHGGGGGGGPSPSSGYGRQDRL*

>OsHAP3H_LOC_Os08g07740.1

MKSRKSYGHLLSPVGSPPLDNESGEAAAAAAAGGGGCGSSAGYVVYGGGGGGDSPAKEQD

RFLPIANVSRIMKRSLPANAKISKESKETVQECVSEFISFVTGEASDKCQREKRKTINGD

DLLWAMTTLGFEAYVGPLKSYLNRYREAEGEKADVLGGAGGAAAARHGEGGCCGGGGGGA

DGVVIDGHYPLAGGLSHSHHGHQQQDGGGDVGLMMGGGDAGVGYNAGAGSTTTAFYAPAA

TAASGNKAYCGGDGSRVMEFEGIGGEEESGGGGGGGERGFAGHLHGVQWFRLKRNTN*

>OsHAP3Hcdo

MKSRKSYGHLLSPVGSPPLDNESGEAAAAAAAGGGGCGSSAGYVVYGGGGGGDSPAKEQDRFLPIANVSRIMKRSLPANAKISKESKETVQECVSEFISFVTGEASDKCQREKRKTINGDDLLWAMTTLGFEAYVGPLKSYLNRYREAEGEKADVLGGAGGAAAARHGEGGCCGGGGGGADGVVIDGHYPLAGGLSHSHHGHQQQDGGGDVGLMMGGGDAGVGYNAGAGSTTTAFYAPAATAASGNKAYCGGDGSRVMEFEGIGGEEESGGGGGGGERGFAGHLHGVQWFRLKRNTN*

>Ghd8_Nipponbare

MKSRKSYGHLLSPVGSPPLDNESGEAAAAAAAGGGGCGSSAGYVVYGGGGGGDSPAKEQD

RFLPIANVSRIMKRSLPANAKISKESKETVQECVSEFISFVTGEASDKCQREKRKTINGD

DLLWAMTTLGFEAYVGPLKSYLNRYREAEGEKADVLGGAGGAAAARHGEGGCCGGGGGGA

DGVVIDGHYPLAGGLSHSHHGHQQQDGGGDVGLMMGGGDAGVGYNAGAGSTTTAFYAPAA

TAASGNKAYCGGDGSRVMEFEGIGGEEESGGGGGGGERGFAGHLHGVQWFRLKRNTN*

>Ghd8_93-11

MKSRKSYGHLLSPVGSPPLDNESGAAAAAAAAGGGGCGSSAGYVVYGGGGGGDSPAKEQD

RFLPIANVSRIMKRSLPANAKISKEAKETVQECVSEFISFVTGEASDKCQREKRKTINGD

DLLWAMTTLGFEAYVGPLKSYLNRYREAEGEKADVLGGAGGAAAARHGEGGCCGGGGGA

DGVVIDGHYPLAGGLSHSHHGHQQQDGGGDVGLMMGGGDAGVGYNAGAGSTTTAFYAPAA

TAASGNKAYCGGDGSRVMEFEGIGGEEESGGGGGGGERGFAGHLHGVQWFRLKRSTN*

>OsCAR8_Hoshihikari

MKSRKSYGHLLSPVGSPPLDNESGEAAAAAAAGGGGCGSSAGYVVYGGGGGGDSPAKEQD

RFLPIANVSRIMKRSLPANAKISKESKETVQECVSEFISFVTGEASDKCQREKRKTINGD

DLLWAMTTLGFEAYVGPLKSYLNRYREAEGEKADVLGGAGGAAAARHGEGGCCGGGGGGA

DGVVIDGHYPLAGGLSHSHHGHQQQDGGGDVGLMMGGGDAGVGYNAGAGSTTTAFYAPAA

TAASGNKAYCGGDGSRVMEFEGIGGEEESGGGGGGGERGFAGHLHGVQWFRLKRNTN*

>AtNFYB5_AT2G47810

MAGNYHSFQNPIPRYQNYNFGSSSSNHQHEHDGLVVVVEDQQQEESMMVKEQDRLLPIANVGRIMKNILPANAKVSKEAKETMQECVSEFISFVTGEASDKCHKEKRKTVNGDDICWAMANLGFDDYAAQLKKYLHRYRVLEGEKPNHHGKGGPKSSPDN*

>AtNFYB4_AT1G09030

MTDEDRLLPIANVGRLMKQILPSNAKISKEAKQTVQECATEFISFVTCEASEKCHRENRKTVNGDDIWWALSTLGLDNYADAVGRHLHKYREAERERTEHNKGSNDSGNEKETNTRSDVQNQSTKFIRVVEKGSSSSAR*

>Zm00001d042197

MSHTSNFTGFSQLEHPQPQRNSRASSSTTHDANVRHDNNLLPIANVGRIMKDALPPQAKISKHAKETIQECTTEFVGFVTGEASERCRRERRKTINGDDICHAMRSLGLDHYADAMRRYLQRYRETEELAAALNSGGGGHDGNAIQIDVRDELSIFKGSNQQGGRD*

>ZmNFYB1_Zm00001d042196

MADHHHHHHHGHPPDGPGGAGDQLEVIKEQDRLLPIANVGRIMKQILPPNAKISKEAKET

MQECVSEFISFVTGEASDKCHKEKRKTVNGDDVCCAFGALGFDDYVDPMRRYLHKYRELE

GDRAASSRGGGGGPAGAADPASASAAAGPSPSAASAGHFMFGAAMDRPDNNSSAGARPF

>AtNFYB6_[AT5G47670](http://www.ensemblgenomes.org/id/AT5G47670)

MERGGFHGYRKLSVNNTTPSPPGLAANFLMAEGSMRPPEFNQPNKTSNGGEEECTVREQD

RFMPIANVIRIMRRILPAHAKISDDSKETIQECVSEYISFITGEANERCQREQRKTITAE

DVLWAMSKLGFDDYIEPLTLYLHRYRELEGERGVSCSAGSVSMTNGLVVKRPNGTMTEYG

AYGPVPGIHMAQYHYRHQNGFVFSGNEPNSKMSGSSSGASGARVEVFPTQQHKY

>AtNFYB9_AT1G21970

MERGAPFSHYQLPKSISELNLDQHSNNPTPMTSSVVVAGAGDKNNGIVVQQQPPCVAREQ

DQYMPIANVIRIMRKTLPSHAKISDDAKETIQECVSEYISFVTGEANERCQREQRKTITA

EDILWAMSKLGFDNYVDPLTVFINRYREIETDRGSALRGEPPSLRQTYGGNGIGFHGPSH

GLPPPGPYGYGMLDQSMVMGGGRYYQNGSSGQDESSVGGGSSSSINGMPAFDHYGQYK

>Zm00001eb189490

MDSSFLPAGADNGSAGGANNGGGAAQQAPPIREQDRLMPIANVIRIMRRVLPAHAKISDD

AKETIQECVSEYISFITGEANERCQREQRKTITAEDVLWAMSRLGFDDYVEPLSVYLHRY

REFEGEARGVGLAPAPPRGDHHHHHHSVPPSMLNKSRGPGSGAVMLPHHHHHDMHASMYG

GAVPPPPHHGFLMPHPQGGHYLPYPYEPTSYGGEHALASGYYGGAAYAPGNNGGSGDGSG

GSASHAPPGGSGGGFDHPHTFAYK

>Zm00001eb253260

MDSSSFLPAAGAENGSAAGGANNGGAAQQHAAPAIREQDRLMPIANVIRIMRRVLPAHAK

ISDDAKETIQECVSEYISFITGEANERCQREQRKTITAEDVLWAMSRLGFDDYVEPLGAY

LHRYREFEGDARGVGLVPGAAPSRGGDHHPHSMSPAAMLKSRGPVSGAAMLPHHHHHHDM

QMHAAMYGGTAVPPPAGPPHHGGFLMPHPQGSSHYLPYAYEPTYGGEHAMAAYYGGAAYA

PGNGGSGDGSGSGGGGGSASHTPQGSGGLEHPHPFAYK

>AtNFYB3_AT4G14540

MADSDNDSGGHKDGGNASTREQDRFLPIANVSRIMKKALPANAKISKDAKETVQECVSEF

ISFITGEASDKCQREKRKTINGDDLLWAMTTLGFEDYVEPLKVYLQKYREVEGEKTTTAG

RQGDKEGGGGGGGAGSGSGGAPMYGGGMVTTMGHQFSHHFS

>AtNFYB2_AT5G47640

MGDSDRDSGGGQNGNNQNGQSSLSPREQDRFLPIANVSRIMKKALPANAKISKDAKETMQ

ECVSEFISFVTGEASDKCQKEKRKTINGDDLLWAMTTLGFEDYVEPLKVYLQRFREIEGE

RTGLGRPQTGGEVGEHQRDAVGDGGGFYGGGGGMQYHQHHQFLHQQNHMYGATGGGSDSG

GGAASGRTRT

>Zm00001eb108760

MPDSDNDESGGPSNADFSSPREQDRFLPIANVSRIMKKALPANAKISKDGKETVQECVSE

FISFITGEASDKCQREKRKTINGDDLLWAMTTLGFEDYVEPLKLYLHKFRELEGDKAAAG

SQPPPPPSSTHNGAGVPVGYGMYGAGGGSGMIMMMGQPMYPPAASSGYSQQPPHHQMSMG

GKGGAYGHCDGSSSSPSGLRRHDGL

>Zm00001eb327140

MPDSDNESGGPSNAEFSSPREQDRFLPIANVSRIMKKALPANAKISKDAKETVQECVSEF

ISFITGEASDKCQREKRKTINGDDLLWAMTTLGFEDYVEPLKLYLHKFRELEGEKAATTS

ASSGPQPPLHRETTPSSSTHNGAGGPVGGYGMYGGAGGGSGMIMMMGQPMYGGSPPAASS

GSYPHHQMAMGGKGGAYGYGGGSSSSPSGLGR

>AtNFYB10-AT3G53340

MAESQTGGGGGGSHESGGDQSPRSLNVREQDRFLPIANISRIMKRGLPLNGKIAKDAKET

MQECVSEFISFVTSEASDKCQREKRKTINGDDLLWAMATLGFEDYIDPLKVYLMRYREME

GDTKGSGKGGESSAKRDGQPSQVSQFSQVPQQGSFSQGPYGNSQGSNMMVQMPGTE

>AtNFYB8_AT2G37060

MAESQAKSPGGCGSHESGGDQSPRSLHVREQDRFLPIANISRIMKRGLPANGKIAKDAKE

IVQECVSEFISFVTSEASDKCQREKRKTINGDDLLWAMATLGFEDYMEPLKVYLMRYREM

EGDTKGSAKGGDPNAKKDGQSSQNGQFSQLAHQGPYGNSQAQQHMMVPMPGTD

>AtNFYB1_AT2G38880

MADTPSSPAGDGGESGGSVREQDRYLPIANISRIMKKALPPNGKIGKDAKDTVQECVSEF

ISFITSEASDKCQKEKRKTVNGDDLLWAMATLGFEDYLEPLKIYLARYRELEGDNKGSGK

SGDGSNRDAGGGVSGEEMPSW

>Zm00001eb148850

MADDGGSHEGSGGGGGVREQDRFLPIANISRIMKKAVPANGKIAKDAKETLQECVSEFIS

FVTSEASDKCQKEKRKTINGDDLLWAMATLGFEEYVEPLKIYLQKYKEMEGDSKLSTKAG

EGSVKKDAISPHGGTSSSSNQLVQHGVYNQGMGYMQPQLGKSLHCLPCVLAENRRHLQRV

LISAMAASQV

>Zm00001eb289860

MADAPASPGGGGGSHESGSPRGGGGGGGGSVREQDRFLPIANISRIMKKAIPANGKIAKD

AKETVQECVSEFISFITSEASDKCQREKRKTINGDDLLWAMATLGFEDYIEPLKVYLQKY

REGDSKLTSKSSDGSIKKDALGHVGASSSAVQGMGQQGTYNQGMGYMQPQISTLIYCKFA

NQFTAFA

>Zm00001eb065670

MRRAVTENGKIARDARESIQECVSEFISFITSEASDKCVKERRKTINDDDIIWSLGTLGF

EEYVEPLKIYLNNYQEGDIKGSKSSDQNGKK

>Zm00001eb025330

MAASRVSACRRWRPTPAAGARSRTDSSLSPTSGASCAARASDKCVKERRKTINDDDIIWS

LGTLGFEEYVEPLKIYLNNYRE

>Zm00001eb125090

MSEVEANTGGGGKEQDRFLPVANIGLIMRRAVPENGKIARDARESIQECVSEFISFITSE

ASDKCVKERRKTINDNDIIWSLGTLGFEEYVEPLKIYLNNYRESILFGGNVKGHGHTLEV

DSKSKRCPLASQQEEHKS

>Zm00001eb344280

MSEVEANAGGRGKEQDRFLPIANIGRIMRRAVPENGKIARDARESIQECVSEFISFITSE

ASDKCVKERRKTINGDDIIWSLGTLGFEEYVEPLKIYLNNYQEGDTKGSKSSDQNGKKQM

LLNGELGSS

>Zm00001eb253270

MGRKAKRCAGKKDDGREEKVAAAPAAEGASSSSSDGEGAGTAATGLPMANLVRLMRQVIP

KGVKISSRAKDLTHDCAVEFVGFLAGEASELARAQHRRTISPEDFTRSLQALGFDDYVRP

MSTYISRYREQATSPAGYSGGFARRPPPRATAAPCVSDEETRSRVPPHGEHGDGWRTSVN

TPAPGGHGYDYTSPDNM

>Zm00001eb302780

MSKAQGSNDHQHEHEDPEGSKPLEEYTIPKGTITRIMRQVLPQDSRVTGGAKETMDQCIV

QFSTALVRAATQECRRDRRLTITADDLIVGFANLGLADYVQPMSVYLRLYRETVNNQQQA

VAPPSPTVQRGTTTAVPPPPPNLTLQLGLPSVPDVTELARDTDVYALWHGAAPAAGSTSA

SSVAPMPPPAADGDEDE

>Zm00001eb380770

MNNPQNPKASAPCTLPPELPKEAVATDEAPPPMGNNNNTESATATMVREQDRLMPVANVS

RIMRQVLPPYAKISDDAKEVIQECVSEFISFVTGEANERCHTERRKTVTSEDIVWAMSRL

GFDDYVAPLGAFLQRMRDDSDHGGEERGGPAGRGGSRRGSSSLPLHCPQQMHHLHPAVCR

RPHQSVSPAAGYAVRPVPRPMPASGYRMQGGDHRSVGGVAPCSYGGALVQAGGTQHVVGF

HDDEASSSSENPPPEGRAAGSN

>Zm00001eb019920

MPDSDNDSGGPSNAGGELSSPREQDRFLPIANVSRIMKKALPANAKISKDAKETVQECVS

EFISFITGEASDKCQREKRKTINGDDLLWAMTTLGFEDYVEPLKHYLHKFREIEGERAAA

SAGASGSQQQQQQGELPRGAANAAGYAGYGAPGSGGMMMMMMGQPMYGGSQPQQQPPPPQ

PPQQQQQHQQHHMAIGGRGGFGQQGGGGGSSSSSGLGRQDRA

>Zm00001eb172020

MKNRKGYGHQGHLLSPVGSPLSDNESGAAAAAGGGGCGSSVGYCGGGGGESPAKEQDRFL

PIANVSRIMKRSLPANAKISKEAKETVQECVSEFISFVTGEASDKCQREKRKTINGDDLL

WAMTTLGFEAYVAPLKSYLNRYREAEGEKAAVLGGGARHGEGGGAADDAGPLAAGGGAGD

GVDRAGHDDDAHVGLMMGASSVGFGSGGGAAPSYYAAASRKAYGAGEGSKVMEFEGEEEN

GGVQRGRGFASHLHGAVQW

>Zm00001eb142030

MAAGHHGQPPDGEDGRRAVVGGEQDRLLPIANVGRIMKQILPPNAKISKEAKETMQECVS

EFIGFVTGEASDKCHKEKRKTVNGDDLCWAFGALGFDDYVDPMRGYLHKYREVEGDRAAA

AASSSRGGGDHHPASASTSTSPAAAAAPGHFMFGAAAIDRPDNNTSSARPF

>Traes_1DL_986875DD3.1
MSEAVGTPESGGAKEQERFLPIANIGRIMRRGVPENGKIAKDAKESIQECVSEFISFITS
EASDKCMKEKRKTINGDDLIWSMGTLGFEDYVEPLKLYLKLYREMEGDTSKGSKSEQAGK
KEVALNGQPGSSFNGM*
>Traes_6AL_477C798AB.1
MGGSSKKRGGQRGEGDAENPAAEGGSALPMANVVRLMRRVLPSNVKIAETAKQLTHDCAV
EFVGFVGGEASERARSEHRRTVAPEDFTWSCQSLGLDSYVQPMQTYLQGYREYDIARGRS
SRGARPPAPPAIASFLPPGQPVTVTEEELEFLRSVVPPPPEGY*
>Traes_3B_924B78EE5.1
PVFSLFPTDERPGSVVFPIAPARAASPSPASDLPFPAPPRSANHRPPPPPPFSPLRSSSR
ARVFIRVSGRGMADDDSGSPRGGGGVREQDRFLPIANISRIMKKAVPANGKIAKDAKETL
QECVSEFISFVTSEASDKCQKEKRKTINGDDLLWAMATLGFEEYVDPLKIYLQKYRDMEG
DSKLTSKSGEGSVKKDIIGAHSGATSSNAQAMVQHGAYAQGMGYMQPQYHNGDT*
>Traes_2DS_21D187BC9.1
LRWAAMPDSDNDSGGPSNADFSSPKEQDRFLPIANVS
>Traes_2AS_B3BC47BB0.1
MKKALPANAKISKDAKETVQECVSEFISFITGEASDKCQREKRKTINGDDLLWAMTTLGF
EDYMEPLKLY
>Traes_1AL_8BC67B52F.1
MSEAVGTPESSGAKEQERFLPIANIGRIMRRGVPENGKIAKDAKESIQECVSEFISFITS
EASDKCMKEKRKTINGDDLIWSMGTLGFEDYVEPLKLYLKLYREMEGDTSKGSKSEQAAK
KVGALNGQPGSSFNGM*
>Traes_6DL_187F01785.1
MGGSSKKRGGQRGEGDGENPAAEGGSALPMANVVRLMRRVLPSNVKIAETAKQLTHDCAV
EFVGFVGGEASERARSEHRRTVAPEDFTWSCQSLGFDSYVQPMETYLRGYREYDIARGRS
SRGARPPAPPATASFLPPGQPVTVTEEELEFLRSVVPPPPEGY*
>Traes_3AL_EF72B5A0D.1
SDKCQKEKRKTINGDDLLWAMATLGFEEYVDPLKIYLQKYRDMEGDSKLTSKSGEGSVKK
DIIGAHSGATSSNAQAMVQHGGYAQGMGYMQPQYHNGDT*
>Traes_3DL_136F5F731.1
MATLGFEEYVDPLKIYLQKYRDMEGDSKLTSKSGEGSVKKDIIGAHSGATSSNAQAMVQH
GAYAQGMGYMQPQYHNGDT*
>Traes_1BL_6F6239DF9.2
MSEAVGTPESGGAKEQERFLPIANIGRIMRRGVPENGKIAKDAKESIQECVSEFISFITS
EASDKCMKEKRKTINGDDLIWSMGTLGFEDYVEPLKLYLKLYREVIFIHSFVPLFLCCLC
YACAACERNSEFGVRRRFGGYSIGVGEL*
>Traes_7DS_A1673EBA1.2
MENADVPNGATAPAPTQATPVVREQDRLMPIANVIRIMRRALPAHAKISDDAKEAIQECV
SEFISFVTGEANERCHMEHRKTVNAEDIVWALNRLGFDDYVVPLSVFLHRMRDPEAATGG
AAAGDRRAVTSAPPRAAPPVLHAMPLHAQMQRPMYAPPALVQVQNQMQRPIYAPPAPMQV
QNQVQWPMYAPPPPVQVQMQRGVYAPQAPVQGYAVGITPVRANVDGQCQVFGGERAVAQQ
YYGYGYGEGSSNGGACADEESSSNSVPVPGKGTGEPEPEPAAEESQGKPIQSG*
>Traes_3B_4D37853E1.1
MKQVLPPNAKVSKEAKETMQECVSEFISFVTGEASDKCHKEKRKTVNGDDVCWAFSALGF
DDYVDPMRRYLLKFRELEGDRAAAAASSRGGLPVPDASTSGAGASGSGNFMFEAMDRRDN
TGPGTGRQF*
>Traes_3DS_833AD0EE2.1
MKKALPANAKISKDAKETVQECVSEFISFITGEASDKCQREKRKTINGDDLLWAMTTLGF
EDYVDPLKHYLHKFREIEGERAAATSTSTTPDMPRNNNNA
>Traes_3AS_9151583B9.1
MPESDNDSGGPSNTGGEGELSSPREQDRFLPIANVSRIMKKALPANAKISKDAKETVQEC
VSEFISFITGEASDKCQREKRKTINGDDLLWAMTTLGFEDYVDPLKHYLHKFREIEGERA
AATSTSTSTTPDMPRNNNNNATGYADAPGGMMMMGQPMYGSXPPQQQQQHQHQIAMGGRA
GFPYLGGGGGSSSSSGFGRKE*
>Traes_6DL_A05C1BF04.1
MRRVLPPHAKISDDAKETIQECVSEYISFITGEANERCQREQRKTITAEDVLWAMSRLGF
DDYVEPLSIYLHRFREFEGEARGAGAGHHHHGMPPMMLKPRGAPGPMVPHGDMQMHAAGM
YGGGAMPPHPHHPFHMPPHHGQYPQYEMYGGEHGMAAYYGGPYAPGNGGNNGDGSGSSGN
GHGGDSTPPAGNFEHHQTFGYK*
>Traes_3AS_102EBF3DD.3
RRASDKCVKEKRKTINGDDLIWSMGTLGFEDYVEPLKLHLKLYQQAFEFKRLLLYYNNVH
DQQVLLMSSISYLAIVSAIFTSADMCSGFQFLHRHG*
>Traes_7AS_067BEC7B5.1
MENDGVPNGPAGPAPTQATPVVREQDRLMPIANVIRIMRRALPAHAKISDEAKEAIQECV
SEFISFDTGEANERCRMQRRKTVNAEDIVWALNRLGFDDYVVPLSVFLERMRDPEAGTGG
AAAGYSRAVMSAPPRAASPVIHAVPLQAQRPMYATPAPVQVHNQMQRPVYAPPAPLQVQM
QQGIYGPRAPVHGYAVGMAPVRANVGG*
>Traes_6AL_857D5AAEB.1
MDMGFPGAANGAAAAAAGNGGGGGQAPAVREQDRLMPIANVIRIMRRVLPPHAKISDDAK
ETIQECVSEYISFITGEANERCQREQRKTITAEDVLWAMSRLGFDDYVEPLSIYLHRFRE
FEGEARGAGAGHHHHGMPPMMLKPRGAPGPMVPHGDMQMHAAGMYGGGAMPPHPHHPFHM
PPHHGQYPQYEMYGGEHGMAAYYGGPYAPGNGGNNGDGSGSSGNGHGGESTPPGGNFEHH
QTFGYK*
>Traes_3B_DBC77FFF1.1
MTNREDFIHFSGFTQQPGRLSLPRAPSTSGSSSGDPNGQEGLLPIANVGRIMKDVLPPEA
KVSKHAKEVIQECATEFIGFVTGEASERCRRERRKTVNGDDICHAMTTLGLDNYAGAMRR
YLQRYREGEELAAVLNNHSRSPAPPAPGDGMIQIDVWGELSNSTGNEKHGRD*
>Traes_2AL_AE22E725E.3
MEIAPLDTSSTHPLTRLLPVLLLLLLLASLDIFFSRRGAASVDMKSRKSYGQQQSHLLSP
VGSPSSDNGGGDSPAKEQDRFLPIANVSRIMKRSLPANAKISKEAKETVQECVSEFISFV
TGEASDKCQREKRKTINGDDLLWAMTTLGFEVYVAPLKAYLNRYREVEGEKAAVVGGSRL
GDDDAHSSLSAAGDALAPQYTHGAGDRGVQDGDLGGHDAHVGLMMGVNMGFNPGTGTTFY
AAPGAAHGRRAYGGGEGARGIDFEAAFGGDRGKNGAGGEREFAGHLHGAVQW*
>Traes_2BL_3237AA694.1
MKSRKSYGQQQSHLLSPVGSPSSDNGGGDSPAKEQDRFLPIANVSRIMKRSLPANAKISK
EAKETVQECVSEFISFVTGEASDKCQREKRKTINGDDLLWAMTTLGFEVYVAPLKAYLNR
YREVEGEKAAVVGGSRHGDDDAHSSLSAAGDALAPQYPHGAGDRGVQDGDVGGHDAHVGL
MMGVNMGFSPGTGTTFYAAPGAAHGRRAYGGGEGARGIDFEGAFGGDRGKNGVGGEREFA
GHLHGAVQW*
>Traes_7DS_A07050CB8.1
MRRALPAHAKISDDAKEAIQECVSEFISFVTGEANERCRMQHRKTVNAEDIVWALNRLGF
DDYVVPLSVFLHRMRDPEAGT
>Traes_2BS_F23D9EA93.1
MPDSDNDSGGPSNADFSSPKEQDRFLPIANVSRIMKKALPANAKISKDAKETVQECVSEF
ISFITGEASDKCQREKRKTINGDDLLWAMTTLGFEDYMEPLKLYLHKFRELEGEKAVGAG
GVGALPSPGGSGSQQRESTPRNNGGGGEAGGYGGMYGGAGAGGGGGGMFMMMGQPMYGSP
PAAGYQHPQHHHQMMTGGQGGYGYGDAGAGGGSSSSSGFGRQDRA*
>Traes_7BS_DBAD99848.1
MENADVPNGAAAPAPTQATPVVREQDRLMPIANVIRIMRRALPAHAKISDDAKEAIQECV
SEFISFVTGEANERCHMEHRKTVNAEDIVWALNRLGFDDYVVPLSVFLHRMRDPEAATGA
AXXXXXXXXXXXXXXXXXXXXXXXXXXXXXXXXXXXXRVQVQNQMQRPVYASPAPMQVQN
QVQRPMYAPPAPVQVQMQRGV
>Traes_2DL_DA577AF57.1
MKSRKSYGQQQSHLLSPVGSPSSDNGGGDSPAKEQDRFLPIANVSRIMKRSLPANAKISK
EAKETVQECVSEFISFVTGEASDKCQREKRKTINGDDLLWAMTTLGFEVYVAPLKAYLNR
YREVEGEKAAVVGGSRHGDDDAHSSLSAAGDALAPPYPHGAGDRGVQDSDVGGHDAHVGL
MMGVNMGFSPGTGTTFYAAPGAAHGRRAYGGVEGARGVDFEAAFGGDRGKNGAGGEREFA
GHLHGAVQW*

>Chlamydomonas_reinhardtii_A0A0I9QPW6

MSGDEGDGRDGNSNAREQDRFLPIANISRIMKKALPNNAKIAKDAKETVQECVSEFISFI

TSEASDKCQREKRKTINGDDLLWAMTTLGFEEYLEPLKLYLAKFREAEAATSNKPGGGSG

ANAEAKREAAAAAAAAAAAAAAVSQQQAAQQQMAAQLQAGMAFPGLMPAQFQGLPPGMIP

AGFPGLPLPPGVPGLMMPGGVVPKQEPPK
